# Identification of HIV Tat and NF-κB binding proteins associated with semen-derived extracellular vesicles

**DOI:** 10.1101/2025.04.25.650599

**Authors:** Bryson C. Okeoma, Hussein Kaddour, Wasifa Naushad, Victor Paromov, Ashok Chaudhary, Alessio Noghero, Jack T. Stapleton, Chioma M. Okeoma

**Author notes:** Regeneron Pharmaceuticals, Inc., Tarrytown, NY 10591, USA. These authors contributed equally to this work (Bryson C. Okeoma, Hussein Kaddour, Wasifa Naushad).

## Abstract

Semen-derived extracellular vesicles (SEVs) have been shown to inhibit transactivation of the long terminal repeat (LTR) in human immunodeficiency virus type 1 (HIV-1, or HIV) and, hence, viral replication by blocking the interaction of the virus’s transcriptional activator Tat and host transcription factors NF-κB and Sp1. The ability of SEVs to regulate the activities of transcription factors suggests that SEVs may contain transcription activators and repressors. Here, we identified host proteins in human SEVs that interacted with the Tat and NF-κB subunit p65. Integrative network and pathway enrichment analyses of these complexes revealed associations with an array of biological functions regulating genome transcription. In particular, several proteins in SEVs could bind to both Tat and NF-κB: the scaffolding and cell signaling regulatory protein AKAP9, the G protein signaling regulator ARHGEF28, the small nuclear RNA processor INTS1, the epigenetic reader BRD2, and the transcription elongation inhibitor NELFB. NF-κB p65–bound NELFB also interacted with HEXIM1, another transcription elongation inhibitor, suggesting that SEVs may inhibit HIV propagation through networks of transcriptional regulation and repression.

**One Sentence Summary:** Proteins in vesicles shed from human semen may repress HIV by targeting transcription factors.

## INTRODUCTION

The transcription of human immunodeficiency virus (HIV) involves complex interaction between host cell transcription factors and the viral long terminal repeat (LTR) promoter, which is divided into four functional regions: modulatory element, enhancer, promoter, and Tat-activating region (TAR). The LTR regulatory elements interact with transcription factors (constitutive and inducible), leading to the assembly of a stable transcription complex that stimulates transcription by RNA polymerase II (Pol II). However, in the absence of the virally encoded *trans*-activator protein Tat, which stimulates HIV transcription elongation, the majority of Pol II directed transcripts terminate prematurely (*1, 2*), affecting progeny production. Thus, identifying factors that impair or block the functions of Tat may block provirus transcription and inhibit viral replication. Additionally, inhibiting the function of Tat may also affect other processes in the HIV lifecycle, because it has been shown that Tat also plays a role in HIV reverse transcription (*3, 4*). It is noteworthy that Tat activates HIV transcription through increasing host Pol II processivity through interactions with the cellular target positive transcription elongation factor b (P-TEFb) amongst other factors (*5, 6*). Hence, Tat participates in a positive feedback mechanism that maintains high levels of proviral transcription in HIV-infected cells. Previously, we showed that extracellular vesicles (EVs) derived from human semen (SEVs) inhibit HIV infection (*7–11*) in part by inhibiting HIV transcription through the disruption of the activities of host (NF-κB, Sp1) and viral (Tat) transcription factors (*12*). This significant observation in SEVs has opened a new focus in identifying the factors in SEVs that are responsible for the observed effects.

EVs are released by all cell types and they carry markers of the producer cells. Thus, if the producer cells are healthy or pathologic, EVs may carry markers corresponding to that cellular state (*7, 13*). Importantly, EV-associated cargos are transferred to proximal and distal cells, whereby the cargo reprograms recipient cells and may alter cellular function (*12–16*). The mechanistic details of how EVs achieve their diverse roles are not fully understood; but it is known that nucleic acid-binding proteins and transcription factors are present in EVs (*17*). These molecules may affect EV-directed phenotypic and functional changes in recipient cells. In regards to HIV, EVs play both pro- and anti-viral roles (*13*). For example, EVs from uninfected cell lines, may activate transcription of latent HIV (*18*) or may stimulate production of pro-inflammatory cytokines through HIV TAR RNA (*15*). On the other hand, EVs from *in vivo* sources have been shown to inhibit HIV replication. Our group has reported that SEVs (but not blood-derived EVs), reduce HIV proviral DNA and viral RNA levels in various cell lines and across multiple concentrations of both HIV type 1 and type 2 (*11*). The inhibitory effects of SEVs occur at viral post-entry steps (*11*). Although the exact mechanism(s) by which SEVs inhibit HIV infection is not fully understood, we observed that multiple steps in the HIV lifecycle, including reverse transcription, copying of integrated proviral DNA, and transcription of viral RNA (*11, 12*) are impaired by SEVs. These observations strongly suggest that SEVs inhibit HIV replication through multiple mechanisms (*19*).

One potential mechanism of SEVs-mediated HIV inhibition we previously proposed is the ability of SEVs to inhibit the DNA binding activities of host Sp1 and NF-κB and to block the recruitment of these transcription factors and Pol II to the HIV LTR (*12*). Notably, SEVs inhibit the interactions of Tat with p65 and of Tat with Sp1, without altering host gene expression (*12*). In this manuscript, we aimed to identify the interactomes of Tat and NF-κB p65 in SEVs, which may provide a molecular basis to develop SEVs cargo as novel anti-HIV therapeutics.

## RESULTS

### EVs are present in the semen of men irrespective of their HIV infection (viremic or aviremic) status

EVs were isolated from semen collected from uninfected subjects (HIV-; n=15) and from those who were HIV type 1–infected and viremic, not on antiretrovirals (HIV+ ART-; n=9) or aviremic on antiretrovirals (HIV+ ART+; n=15). The viremic (HIV+ART-) donors were difficult to come by, because the majority of infected individuals are placed on ART upon diagnosis. There were some differences and similarities amongst the SEVs with regard to size, concentration, ζ-potential (mV), and total protein (**Fig. 1A**). There were statistically significant differences in the total protein content of SEVs of uninfected subjects (HIV-ART-, ~1.8 μg/μL) and those infected who are not on ART (HIV+ ART-, ~0.8 μg/μL), or on ART (HIV+ ART+, ~1.4 μg/μL) (**Fig. 1A**). TEM analysis showed that SEVs of uninfected subjects, HIV+ ART-, and HIV+ ART+ SEVs have similar morphology and are heterogenous in size and electron density (**Fig. 1B**). Furthermore, based on the system generated standards using the manufacturer’s protocol of the Jess™ Simple Western system, an automated capillary-based assay for immunodetection of EV markers. (**figs. S1, A and B**), we observed that the isolated SEVs were positive for the typical EV markers CD9, CD63, CD81, and Flotillin-1 (**Fig. 1C, and figs. S1, C to F**). Because differences exist in the total protein content of the SEVs (**Fig. 1A**), we assessed the protein footprints of the SEVs using the Jess™ Simple Western system and showed that the protein footprints are similar irrespective of whether equal protein weight (1.5 µg) or equal particle number (5×10^7^) was used (**Fig. 1D**).

**Figure 1.**
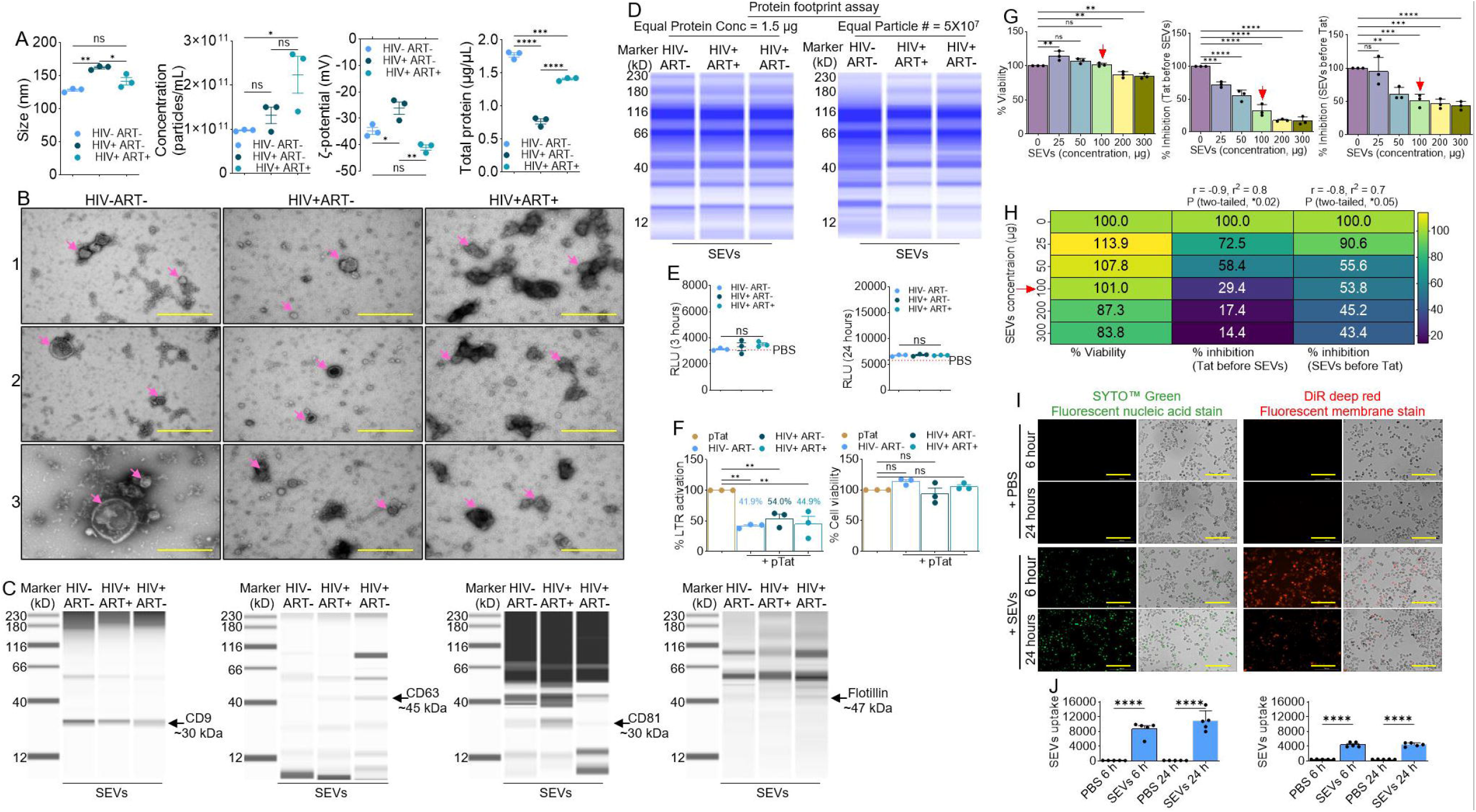
SEVs do not transactivate HIV promoter but inhibit Tat-dependent HIV LTR transactivation: **A**) Characterization of the properties (particle size, concentration, ζ-potential) and total protein of SEVs from uninfected subjects (HIV-ART-), HIV+ART-, HIV+ART+ donors. **B**) Representative TEM images of the morphology of uninfected subjects (HIV-ART-), HIV+ART-, HIV+ART+ SEVs. **C**) Image of lane views of markers (CD6, CD63, CD81, Flotillin) of EVs analyzed by capillary-based western assay using the JESS Simple Western. **D**) Chemiluminescent total protein detected using the JESS Simple Western. **E**) Effects of PBS or 100 μg/mL uninfected subjects (HIV-ART-), HIV+ART-, HIV+ART+ SEVs on HIV LTR promoter activity assessed in TZM-bl indicator cells for 3 hours (left) and 24 hours (right). **F)** Effects of PBS or 100 μg/mL uninfected subjects (HIV-ART-), HIV+ART-, HIV+ART+ SEVs on Tat-mediated HIV LTR promoter transactivation (left) and cell viability (right) assessed in TZM-bl indicator cells. The numbers above the top bars denote % SEVs-mediated inhibition of Tat-directed HIV promoter transactivation. **G**) SEV concentration dependent effects on viability (left) and time-of-SEVs addition on inhibition of Tat-mediated LTR promoter transactivation for co-treatment of cells with Tat and SEVs (middle) and pre-treatment of cells with SEVs for 6 hours prior to the addition of Tat (right). **H**) Heat map of cell viability and time-of-SEVs addition on inhibition of Tat-mediated LTR transactivation for co-treatment and pre-treatment. **I**) Representative 10 X images of TZM-bl cells incubated with SEVs labelled with SYTO™ Green Fluorescent nucleic acid stain (green fluorescence, left) and DiR deep red Fluorescent membrane stain (red fluorescence, right) taken at 24 hours using automated Lionheart FX microscope (Biotek). **J**) Uptake of labeled SEVs by TZM-bl cells analyzed by quantifying 3 fields of view with Gen5 software and presented as raw relative florescence units (RFU). All experiments were repeated three times. Statistical differences were assessed by ordinary one-way ANOVA and unpaired t test. **** p < 0.0001, *** p < 0.001, ** p <0.01, * p < 0.05, ns = non-significant. Pooled donor semen samples used were as follows: Uninfected subjects (HIV-ART-) (n=15, split into 3 groups of 5/group), HIV+ART- (n=9, split into 3 groups of 3/group), HIV+ART+ (n=15, split into 3 groups of 5/group). Panel B scale bar: 500 nm. Panel I scale bar: 200 µm. Experiment was repeated three times.

### SEVs do not alter basal HIV LTR activity, irrespective of whether the SEV donors were HIV viremic or aviremic

Previously, Welch *et al*., showed that SEVs isolated from uninfected subjects did not activate HIV promoter (*12*). However, whether SEVs from HIV+ART- or HIV+ART+ will activate HIV LTR remains to be determined. We examined the effect of SEVs from uninfected subjects, HIV+ ART-, and HIV+ ART+ donors and assessed their effect on the basal activity of the HIV promoter. PBS (vehicle) was used as negative control. TZM-bl cells were treated with either 100 µg of each of the SEVs alone or with PBS, and bioluminescence levels were examined 24 hours later. Basal activity of the HIV promoter at 3 hours and at 24 hours did not change in the presence of any of the SEVs compared to PBS (**Fig. 1E**).

### SEVs inhibit Tat-dependent HIV LTR transactivation irrespective of the donor HIV status (uninfected, viremic, or aviremic)

To examine the effect on Tat-dependent HIV LTR activation of SEVs from uninfected subjects, HIV+ ART-, and HIV+ ART+, TZM-bl cells were transfected with a Tat expression vector (pCMV Tat, hereafter referred to as pTat) (*20*) followed by treatment with SEVs (100 µg/ml). SEVs from uninfected subjects, HIV+ ART-, and HIV+ ART+ inhibited Tat-dependent transcription of a luciferase reporter by approximately half, with no differences in cell viability as measured by MTT assay (**Fig. 1F**). Because there was little difference in SEV-mediated inhibition of Tat between the groups, for subsequent experiments, we used only SEVs from uninfected subjects, hereafter referred to as SEVs.

### Time of addition study shows that SEVs are a bona fide inhibitor of HIV Tat

Here, we explored the effect of time-of-SEVs addition on inhibition of Tat-mediated LTR transactivation. First, increasing concentrations (25 to 300 µg) of SEVs were added to equivalent numbers of TZM-bl cells that were then cultured for 24 hours. MTT assay revealed that 200 and 300 µg of SEVs reduced TZM-bl cell viability to 87% and 84% respectively, whereas 100 µg of SEVs had no effect on cell viability (**Figs. 1, G and H**). Co-treatment of cells with Tat and SEVs synchronously show that pre-incubation of Tat and SEVs for 1 hour at 37 °C before treatment of cells resulted in a dose-dependent inhibition of Tat-mediated LTR transactivation (**Figs. 1, G and H**). Likewise, pre-treatment of cells with SEVs for 6 hours prior to the addition of Tat also resulted in a dose-dependent inhibition of Tat-mediated LTR transactivation (**Figs. 1, G and H**). Of note, 100 µg of SEVs significantly inhibited Tat-mediated LTR promoter transaction by 29.4% by co-treatment and 53.8% by pre-treatment (**Figs. 1, G and H**).

### SEVs are internalized by TZM-bl indicator cells

Internalization of labeled SEVs by TZM-bl indicator cells were accessed as evidenced by green signal from the cell-permeant nucleic acid label SYTO Green Fluorescent Nucleic Acid Stains (**Fig. 1I**) and lipophilic fluorescent membrane stain DiR deep red (**Fig. 1I**). Uptake of SEVs was quantified at 6 and 24 hours for nucleic acid and membrane stains respectively (**Fig. 1J**).

### Identification of Tat binding proteins and its interactomes in SEVs

We have shown that Tat interacts with cellular transcription factors, such as NF-κB (*12*), however, the identity of SEVs proteins that interact with Tat is yet to be determined. To begin to unravel SEV proteins that interact with Tat, extracts from overnight culture of *E. coli* expressing His-Tat were purified on magnetic beads, to which SEVs extract was added and incubated. The beads were washed with Imidazole-containing buffer to remove non-specific proteins. Specific interactomes of Tat in SEVs were eluted with laemmli/BME solution and subjected to SDS-PAGE (**Fig. 2A**). Controls included *E. coli*, SEV extracts, and Tat alone. Proteins in the 10 to 35 molecular-weight bands representing Tat monomer, dimer, and tetramer were excised in the Tat alone, SEVs alone, and SEVs and Tat interactome lanes (**Fig. 2A, arrows**) were excised and subjected to MS analysis to identify interactomes of Tat in SEVs (**table S1**). Two-way Venn diagrams analysis (**table S2**) identified SEVs proteins that were present in Tat complexes (**Fig. 2A**) in two independent experiments. In the first experiment, 178 proteins were present in SEVs, whereas 60 proteins were present in Tat eluates from SEVs. Of these 60, 35 were unique to the Tat eluates (**Fig. 2B**). Similarly, in the second experiment, 161 proteins were present in SEVs whereas 54 proteins were present in Tat eluates from SEVs. Of these 54, 27 proteins were unique to the Tat eluates (**Fig. 2B**). Integration of the Tat eluate–specific proteins from the two experiments yielded 16 proteins (EVPL, GVIN1, PTBP1, AKAP9, SPTCS, ARHGEF28, PTPRQ, GSTCD, ITSN1, INTS1, LRIQ1, CC157, GNPTA, GPD1L, ANK3, BRD2) that were reproducibly part of Tat interactome in SEVs (**Fig. 2C and table S2**). The enrichment levels of these 16 proteins in Tat eluates from SEVs was visualized by heatmap analysis (**Fig. 2D**). The 16 proteins were similarly enriched in Tat eluates from SEVs in experiments 1 and 2, except for EVPL, PTBP1, and ARG28. The differences in enrichment levels may be related to the donors of the semen used to isolate SEVs.

**Figure 2.**
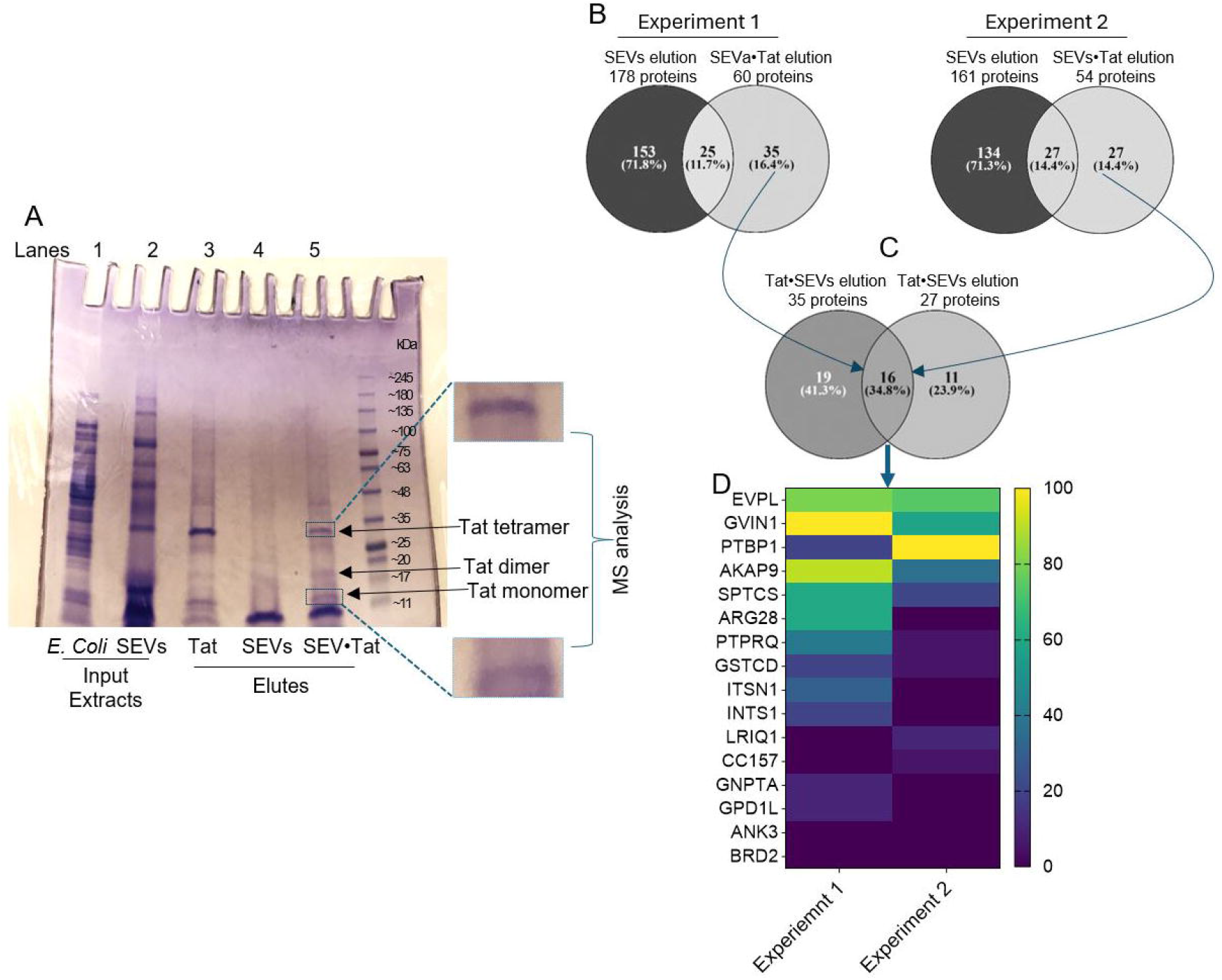
Identification of interactomes of Tat in SEVs: **A**) SDS-PAGE gel showing the interactomes of Tat in SEVs. Controls include *E. coli* and SEVs extracts, or Tat alone. Protein bands (arrows) that were excised (enlarged) were subjected to MS analysis. **B)** 2-way Venn diagram used to identify the interactomes of Tat in SEVs. **C)** 2-way Venn diagram used to identify the interactomes of Tat in SEVs that are common to the two experiments. **D)** Heatmap showing the intensities of the 16 common proteins within the interactomes of Tat in SEVs identified in two separate MS analyses. Pooled uninfected subjects (HIV-ART-) (n=15, split into 2 groups of 7 for group 1 and 8 for group 2).

### Predicted Tat and SEVs network and protein-protein interaction (PPI) clusters identify molecules relevant to HIV transcription

The 16 proteins in the interactomes of Tat in SEVs shown in **Figs. 2C, 2D** are in network with various host proteins as shown on the interactome map in **Fig. 3A**. One of the proteins, polypyrimidine tract binding protein 1 (PTBP1), is a nuclear enzyme with protein and nucleic acid binding properties. On the network, PTBP1 is depicted to directly activate IL-2 and MYC, IL-2 activates MYC, which inhibits BRD2, whereas miR-124 regulates IL-2, PTBP1, and STAT3 (**Fig. 3B**). Detailed relationship type with other molecules is presented in **table S3**. For example, BRD2 and PTBP1 are key proteins present in the interactomes of Tat in SEVs. As depicted, BRD2 interacted with BRD2 and MYC with relationships that includes molecular cleavage, protein-DNA interactions, and regulation of binding, whereas PTBP1 interacted with CSDE1, CTNNB1, GLS2, IL2, MYC, alanine, Hmga2, CTNNB1, HIF1A, HOXA11-AS, MALAT1, DNTT, MYC, NOVA2, PTBP1, with relationships that include causation, expression, protein-DNA, protein-RNA, and protein-protein interactions (**table S3**). Whether the alanine is an independent amino acid or a residue within a protein is unknown. The network map shown in **Fig. 3A** indicates that PTBP1 blocks CSDE1 signaling. CSDE1 signaling is a part of CRD-mediated mRNA stability complex that enables RNA stem-loop binding activity that is involved in IRES-dependent viral translational initiation; nuclear-transcribed mRNA catabolic process, no-go decay; and stress granule assembly.

**Figure 3.**
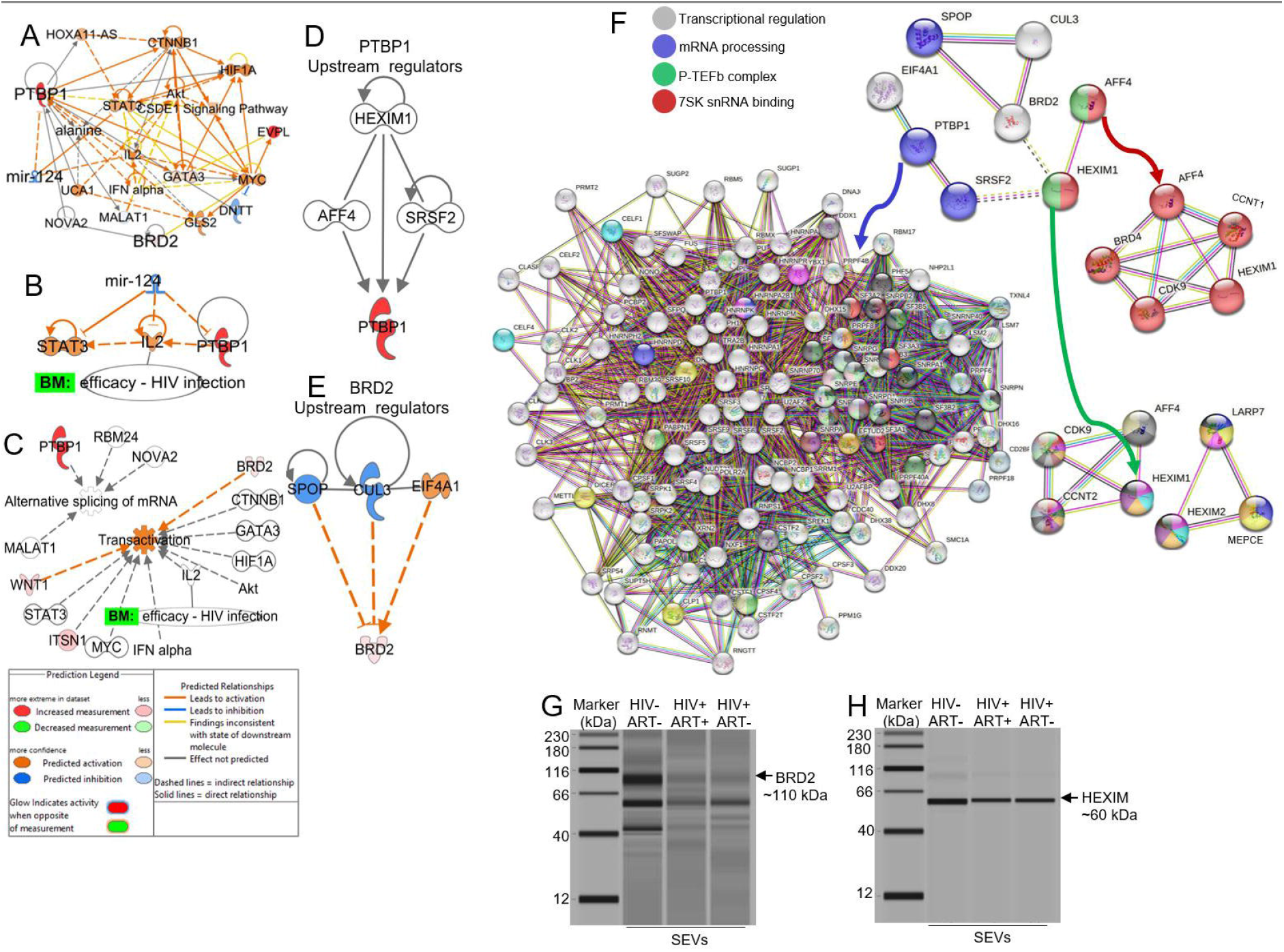
Network of interactomes of Tat in SEVs: **A to E**) Identity of the various host proteins in network with the 16 common proteins in the interactomes of Tat in SEVs and their different interactors. The analysis was conducted with the IPA analysis suite. **F**) PPI of interactomes of Tat in SEVs using the STRING database. **G to H**) Images of lane views of SEVs-associated BRD2 (left) and HEXIM1 (right) analyzed by capillary-based western assay using the JESS Simple Western (ProteinSimple). Pooled donor semen samples used to generate data for network analysis were as follows: Uninfected subjects (HIV-ART-, n=15), HIV+ART- (n=9), HIV+ART+ (n=15).

Further analysis of disease and function, linked PTBT1 (*21*) to alternative splicing of mRNA by spliceosome with Nova2, Malat1, and RBM24 (*22*) as members of the cluster (**Fig. 3C**). Moreover, PTBP1 protein was shown to decrease transcriptional activation of a luciferase reporter gene that is dependent on IFNα (*23*). Likewise, BRD2 which regulates gene transcription (*24*), is in cluster with other proteins known to decrease (CTNNB1, HIF1A) (*25, 26*) or increase or both decrease/increase (MYC, WNT1, ITSN1, Akt, STAT3) (*27–32*) transactivation of genes (**Fig. 3C**). BRD2 is in cluster with IL2, a pro-inflammatory T_Helper_ type 1 cell–associated cytokine that was used by GlaxoSmithKline as a biomarker for measuring the efficacy of abacavir/amprenavir/lamivudine/ritonavir in the treatment of HIV infection. Further analysis identified HEXIM1, AFF4, and SRSF2 as upstream regulators that can affect the expression, transcription, or phosphorylation of PTBP1 (**Fig. 3D**), while SPOP, EIF4A1, CUL3 are upstream regulators of BRD2 (**Figs. 3D, E**). Moreover, PTBP1 regulates the CSDE1 and EIF2 Signaling (**fig. S2**). CSDE1 is part of CRD-mediated mRNA stability complex that enables RNA stem-loop binding activity involved in viral translational initiation; nuclear-transcribed catabolic process, no-go decay; and stress granule assembly.

To further identify proteins sharing the same GO terms in the networks, the STRING database was used for enrichment analysis by integrating publicly available dataset with our dataset to identify PPI associated with PTBP1 and BRD2. The PPI shows that interactome clusters of PTBP1 and BRD2 that interact with HEXIM1 and BRD2 also interact with HEXIM1 via SRSF2. The 4 main interactome clusters include transcriptional regulation (EIF4A1, CUL3, BRD2), mRNA processing (PTBP1, SPOP, SRSF2), P-TEFb complex (HEXIM1, AFF4, CDK9, CCNT2, LARP7, HEXIM2, MEPCE), and 7SK snRNA binding (AFF4, HEXIM1, CDK9, CCNT1, BRD4) clusters (**Fig. 3F**). GO enrichment analysis of our gene set for biological processes, molecular functions, and cellular components that are associated with 7SK snRNA binding, mRNA Processing, and P-TEFb complex was performed. Five top GO processes that were identified include positive regulation of snRNA transcription by RNA polymerase II, 7SK snRNA binding, modification by virus of host mRNA processing, and negative regulation of mRNA polyadenylation (**Table S4**). Reactome and WikiPathways suggest involvement with processes such as, interactions of Tat with host cellular proteins, pausing and recovery of HIV elongation, inhibition of host mRNA processing and RNA silencing, and others (**Table S5**).

The Jess™ Simple Western system was used for immunodetection of specific SEVs-associated proteins using antibodies against BRD2 and HEXIM 1 (**Figs. 3G, H**). Our data indicate that both BRD2 and HEXIM1 are enriched in SEVs from the semen of uninfected subjects compared to HIV+ ART- and HIV+ ART+ SEVs (**Figs. 3F, G, figs. S1G, H**).

### Visualization of the subcellular localizations of Tat binding proteins in SEVs

To further understand the cellular function of the proteins in the mRNA processing, 7SK snRNA binding, and P-TEFb complex GO clusters, we determined their subcellular localizations. Proteins in the 7SK snRNA binding and P-TEFb complex clusters are in the cyclin/CDK positive transcription elongation factor complex, Nucleoplasm, DSIF complex, P-TEFb complex, and NELF complex subcellular localizations/compartments (**table S5**). Notably, the NELF complex [GO:0032021] consists of NELFA, NELFB, NELFCD, NELFE, SUPT5H, CCNT1, CDK9 and other proteins (**Fig. 4A, table S6**). The proteins in the NELF complex may associate with RNP polymerase II to induce transcriptional pausing. To identify overrepresented biological processes, cellular components, and molecular functions in the NELF complex PPI interactome, a STRING-augmented GO enrichment analysis was performed. The GO enrichment analyses highlighted, among other findings, that the NELF complex is associated with GO biological processes that are associated with negative regulation of mRNA polyadenylation, negative regulation of DNA-templated transcription elongation, negative regulation of transcription elongation from RNA polymerase II promoter, positive regulation of viral transcription, and negative regulation of gene expression. Furthermore, the NELF complex is also associated with GO cellular components associated with 7SK snRNA binding, chromatin and RNA binding; and GO molecular functions associated with NELF complex, cyclin/CDK positive transcription elongation factor complex, and transcription elongation factor complex (**table S7**). Interestingly, immunoprecipitation assay showed that Tat binds host NELFB in vitro (**Fig. 4B**).

**Figure 4.**
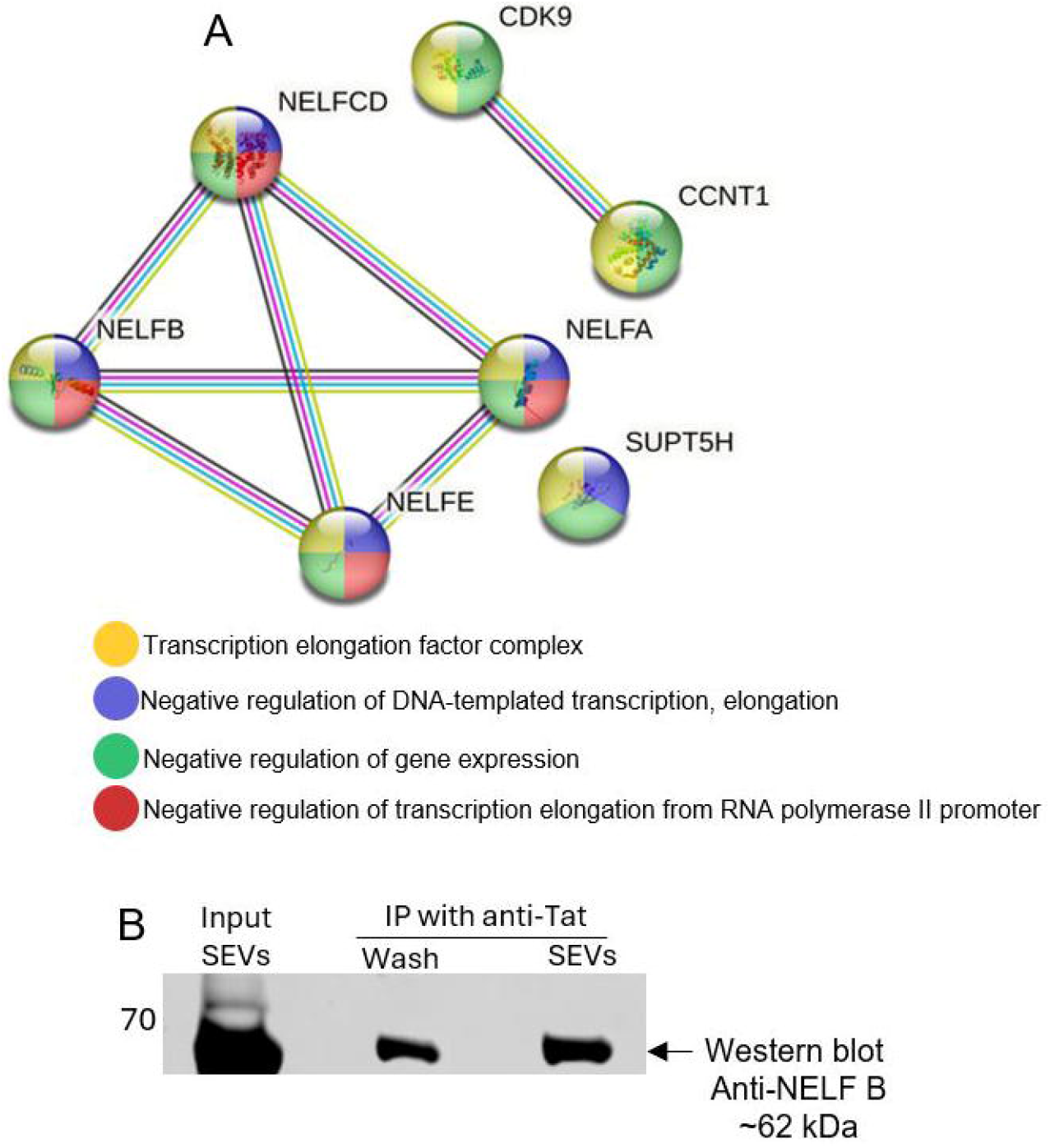
Validation of NELFB as interactomes of Tat in SEVs: **A)** STRING database analysis identifies NELF complex [GO:0032021] consisting of NELFA, NELFB, NELFCD, NELFE and other proteins as interactomes of Tat in SEVs. **B)** Immunoprecipitation and western blot assays showing interaction of SEVs-associated Tat with SEVs-associated NELFB. Experiment was repeated three times. Pooled donor semen samples used to generate data for network analysis were as follows: Uninfected (HIV-ART-, n=15) for panel A and HIV+ART+, n=15 for panel B.

### Identification of interactomes of NF-κB p65 in SEVs

Given that SEVs directly block NF-κB binding to the LTR and inhibits the interactions of Tat with NF-κB (*12*), we sought to identify interactomes of NF-κB p65 in SEVs proteins that may contribute to SEVs-directed inhibition of HIV transcription. We used our previously described EMSA assay (*12*) to show that proteins in SEVs bind NF-κB p65 (**Fig. 5A**). These NF-κB p65 and SEVs complexes were subjected to mass spectrometric (MS) analysis. In total, 83 proteins from SEVs were found in complex with NF-κB p65 (**Fig. 5B**). Of the 83 proteins, BRD2 was present in interactomes of NF-κB p65 in SEVs (**Fig. 5B, blue arrow**). Other bromodomain family of proteins (BRD3, BRDT, orange arrows), along with NELFB (orange arrow), which is an essential component of the NELF complex that negatively regulates the elongation of transcription by Pol II and plays a critical role in suppressing HIV transcription (*33–41*), are also part of the interactomes of NF-κB p65 in SEVs (**Fig. 5B**). Examination of these proteins based on literature-curated functions and pathway analysis using Molecular Interaction Search Tool (MIST) reveals complex PPI and genetic interactions between SEVs proteins that interact with NF-κB p65 (**Fig. 5C**). Gene Set Enrichment Analysis (GSEA), performed with WebGestalt (WEB-based Gene SeT AnaLysis Toolkit) identified top five processes, including metabolism of proteins, innate immune system, axon guidance, phase II - conjugation of compounds, and RNA polymerase II transcription enriched in the interactome (**Fig. 5D**).

**Figure 5.**
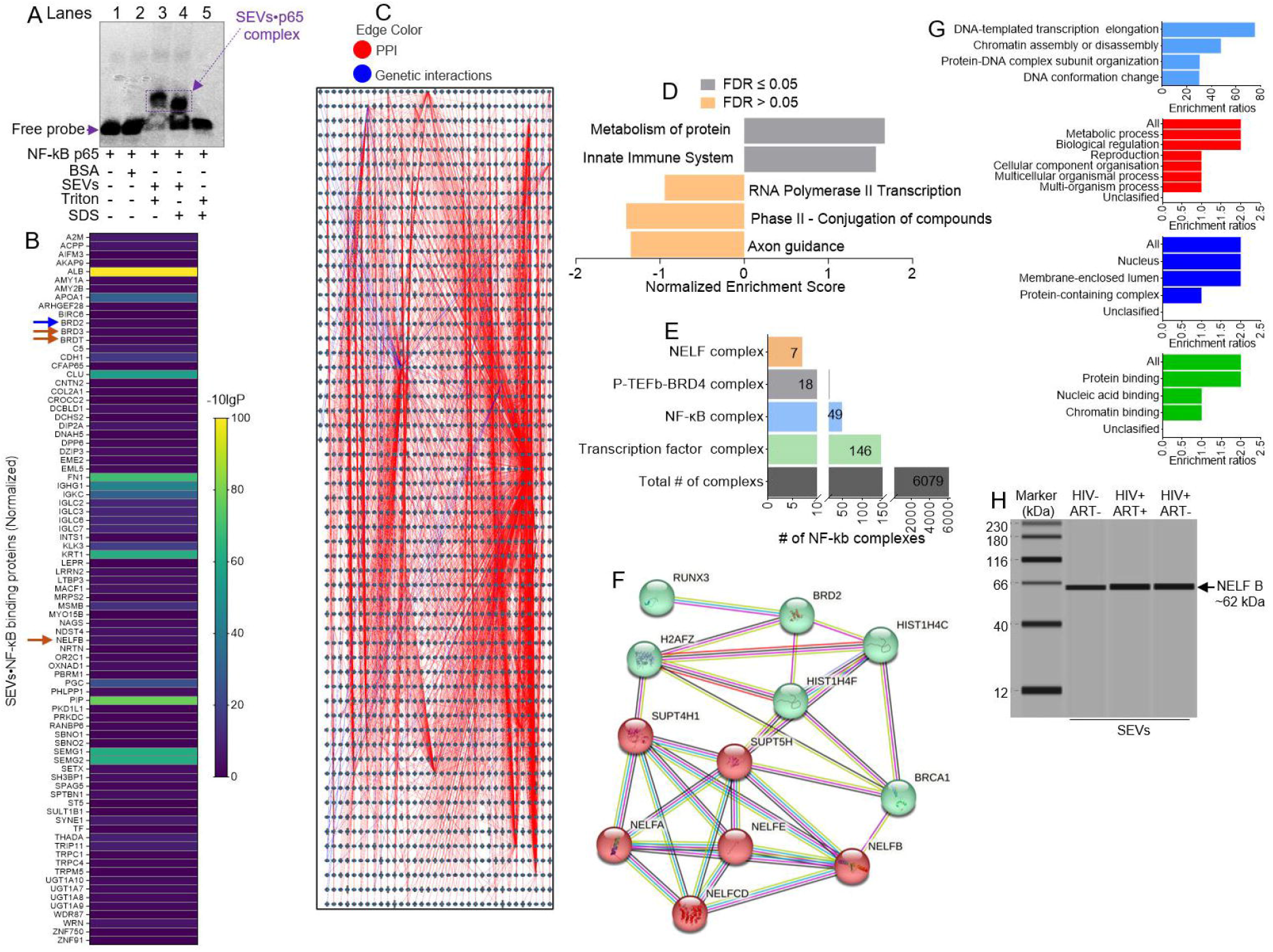
Identification of the Interactomes of NF-κB p65 in SEVs: **A)** The complex formed between proteins in SEVs and NF-kB p65 as identified by EMSA assay. **B)** Heatmap showing the intensities of 83 the interactomes of NF-κB p65 in SEVs, including BRD2 (blue arrow), other bromodomain family of proteins (BRD3, BRDT, orange arrows), and NELFB (orange arrow). **C)** PPI and genetic interactions between the interactomes of NF-κB p65 in SEVs as revealed by MIST functions and pathway analysis. **D)** Top five processes identified by WebGestalt GSEA. **E)** NELFB complexes, including NELF, P-TEFb-BRD4, NF-κB, and transcription factor complexes, with 7, 18, 49, and 146 genes respectively. **F)** PPI analysis showing NELF – BRD2 network clusters. **G)** NELF-BRD2 GSEA, GO biological process, GO cellular component, and GO molecular functions. **H**) Image of lane view of SEVs-associated NELFB analyzed by capillary-based western assay using the JESS Simple Western (ProteinSimple). Pooled donor semen samples used were as follows: Uninfected subjects (HIV-ART-, n=15), HIV+ART- (n=9), HIV+ART+ (n=15).

### NELFB is a part of the interactomes of NF-κB p65 in SEVs

Since NELFB is present in the interactomes of NF-κB p65 in SEVs (**Fig. 5B**), we analyzed NF-KB complexes (**table S8**). Although many complexes (6079) were identified (**table S8 and Fig. 5E**), we found that NELF, P-TEFb-BRD4, NF-κB, and transcription factor complexes have 7, 18, 49, and 146 target genes, respectively (**Fig. 5E**). PPI analysis showed NELF and BRD2 in two network clusters that were linked by SUPT4H1, SUPT5H, and BRCA1 (**Fig. 5F**). Of note, it has been shown that NELF acts with DRB sensitivity-inducing factor (DSIF), a heterodimer of SUPT4H1 and SUPT5H, to cause transcriptional pausing of RNA polymerase II (*42*). GSEA for NELF-BRD2 as well as GO biological process, GO cellular component, and GO molecular functions suggest enrichment of processes related to transcription, chromatin assembly/disassembly, reproduction, protein, nucleic acid, and chromatin binding (**Fig. 5G**). The Jess™ Simple Western system was used for immunodetection of SEVs-NELFB protein using anti NELFB antibody (**Fig. 5H and fig. S1I**). Our data indicated that NELFB is enriched in SEVs from the semen of HIV+ ART- and HIV+ ART+ SEVs compared to uninfected subjects (**Fig. 4D**).

### BRD2 and NELFB are in the same complex in SEVs

We integrated interactomes of Tat and NF-κB p65 in SEVs and identified four proteins, including A-kinase anchoring protein 9 (AKAP9), Rho guanine nucleotide exchange factor 28 (ARHGEF28), bromodomain containing 2 (BRD2), and integrator complex subunit 1 (INTS1) as common proteins in the interactomes of Tat and NF-κB p65 in SEVs (**Fig. 6A**). The level of enrichment of the molecules is similar between interactomes of NF-κB p65 and interactomes of Tat in SEVs, except for AKAP9 (**Fig. 6B**). To identify possible interactions associated with AKAP9, ARHGEF28, BRD2, INTS1, Ingenuity Pathway Analysis (IPA) proposed 4 connecting proteins, including beta-estradiol, ESR1, TP53, and GRHL2 (**Fig. 6C**). Diseases and functions associated with the interactomes of Tat and NF-κB p65 in SEVs include decompaction of heterochromatin and inflammatory response of bone marrow-derived macrophages mediated by BRD2 (**table S9**). Querying functional database (PPI BIOGRID) for Network Topology-based Analysis (NTA) using WebGestalt showed that all four genes (AKAP9, ARHGEF28, BRD2, INTS1) are in a network with enriched GO categories (the top 10) in RNA biology, including RNA splicing, mRNA processing, 3’-UTR-mediated mRNA stabilization, RNA processing, mRNA stabilization, and mRNA metabolic process (**Fig. 6D**). Additionally, one of the top 10 interactors of BRD2 is Tat (**Fig. 6D**). Enriched GO categories (Top 10) associated with the BRD2 interactors include regulation of post-transcriptional gene silencing and others (**table S10**). PPI assay using immunoprecipitation indicate that Tat physically interacted with BRD2, HEXIM 1, and NELFB (**Fig. 6E**).

**Figure 6.**
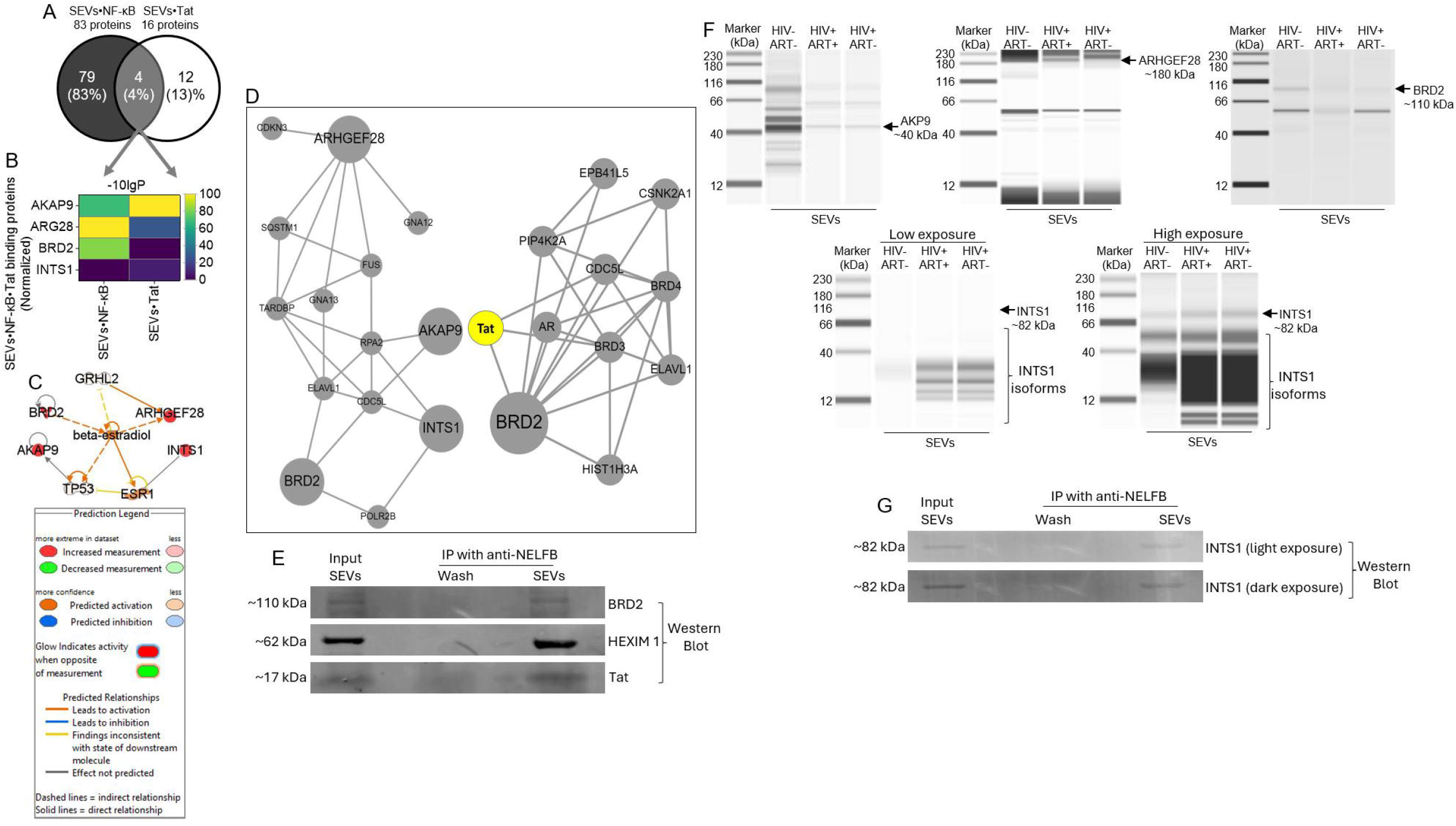
Identification of the interactomes of Tat and NF-κB p65 in SEVs: **A)** Four common proteins (AKAP9, ARHGEF28, BRD2, INTS1) in the interactomes of Tat and NF-κB p65 in SEVs identified by 2-way Venny. **B)** Heatmap depicting the level of enrichment of the 4 proteins that are part of the interactomes of Tat and NF-κB p65 in SEVs. **C)** Other host proteins associated with the interactomes of Tat and NF-κB p65 in SEVs (AKAP9, ARHGEF28, BRD2, INTS1). **D)** WebGestalt NTA of the interactomes of Tat and NF-κB p65 in SEVs (AKAP9, ARHGEF28, BRD2, INTS1) network with top 10 GO enriched categories (left) and showing BRD2 interacting with Tat (right). **E)** Immunoprecipitation and western blot assays showing physical association between SEVs-associated NELFB with BRD2, HEXIM1, and HIV Tat. **F**) Images of lane views of the 4 interactomes of Tat and NF-κB p65 in SEVs (AKAP9, ARHGEF28, BRD2, INTS1) analyzed by capillary-based western assay using the JESS Simple Western (ProteinSimple). **G**) Immunoprecipitation and western blot assays showing physical association between SEVs-associated NELFB with INTS1. Pooled HIV+ART+ donor semen samples (n=15). Experiment was repeated three times.

### Validation of interactomes of Tat and NF-κB p65 in SEVs

The levels of the 4 proteins (AKAP9, ARHGEF28, BRD2, INTS1) present in the interactomes of Tat and NF-κB p65 identified to be enriched in SEVs were analyzed using the Jess™ Simple Western system for validation. The analysis confirmed the presence of AKAP9, ARHGEF28, BRD2, INTS1 (**Fig. 6F and Figs. S1, J to L**). Because NELF complex was shown to interact with the integrator complex subunits (*40*), we used IP and western blot assays to demonstrate physical association of NELFB with INTS1 (**Fig. 6G**). These results corroborate the proteomics data and strongly suggest that AKAP9, ARHGEF28, BRD2, INTS1 are part of the interactomes of Tat and NF-κB p65 in SEVs.

## DISCUSSION

HIV genome encodes viral proteins in a single unspliced (US) polycistronic transcript of 9.2 kb. The host cell spliceosome facilitates alternative splicing leading to the production of over 100 viral RNA (vRNA), including 4kb singly spliced (SS) and 2 kb multiple spliced (MS) vRNA. While SS and US vRNA accumulate in the nucleus, MS vRNA that encodes Rev and Tat are exported to the cytoplasm and translated into Rev and Tat proteins (*2*).

HIV transcription is controlled by upstream LTR, including cis-elements that bind host NF-ᴋB and HIV Tat, which are critical for efficient HIV transcription. Despite its importance, Tat has not been a major focus in terms of its potential for small molecule therapeutics against HIV. In a previous study, we showed that SEVs disrupts NF-ᴋB/Sp1/Tat circuitry to promote transcriptional silencing of HIV (*12*). However, whether SEVs proteins interact with Tat or NF-ᴋB and the interactomes of Tat and NF-κB p65 in SEVs is unknown.

Through a combination of multiple independent assays, we identified Tat and NF-ᴋB binding SEVs-associated proteins as having the potential to inhibit HIV transcription. While SEVs do not alter basal HIV LTR activity, SEVs inhibit Tat-mediated transactivation of HIV LTR, indicating a potential for a beneficial effect to control HIV replication. We found that the cellular nuclear RNA-binding protein, PTBP1, known to associate with HIV RNA (*43, 44*), is part of the interactomes of Tat in SEVs. The observation that PTBP1 is part of the interactomes of Tat in SEVs suggest that PTBP1 may act as a positive or negative regulator for HIV gene expression, RNA stability, and RNA export. Indeed, elevated levels of PTBP1 have been shown to increase cytoplasmic accumulation of HIV RNAs and the release of replication-competent HIV from latently infected cells without inducing cellular stimulation in resting CD4+ T cells (*43*).

We also found that interactomes of Tat in SEVs fall in three clusters: 7SK snRNA binding, mRNA processing, and P-TEFb complex clusters. Proteins in the 7SK snRNA binding cluster are in the P-TEFb complex, DSIF complex, and NELF complex subcellular compartment. Proteins in the mRNA processing cluster are in the U2-type prespliceosome, cleavage body, U4 snRNP, U2 snRNP, and pICln-Sm protein complex. Those in the P-TEFb complex cluster are in the P-TEFb complex, cyclin/CDK positive transcription elongation factor complex, DSIF complex, NELF complex, and Nucleoplasm. These observations are interesting because recruitment of P-TEFb to TAR allows stalled Pol II to transition to transcriptional elongation state (*45*). Thus, Tat and P-TEFb complex plays a critical role in Pol II mediated transcriptional elongation of HIV genes, that involves promoter proximal pausing of newly initiated polymerases by negative elongation factors (DSIF, NELF) and a P-TEFb-mediated transition into productive elongation. These negative elongation factors are modified (DSIF) or replaced (NELF) by positive factors within the SEC (*46*).

Although the primary target of Tat appears to be P-TEFb, Tat also interacts with other cellular cofactors, including Sp1 and NF-kB (*47, 48*) to regulate HIV transcription. While a large body of research is focused on cellular complexes involved in Tat-mediated HIV transcription, our earlier work using EMSA assay was the first to show that factors in SEVs bind NF-ᴋB p65 (*12*). In this current study we identified NELFB and BRD2 as interactomes of NF-ᴋB p65 in SEVs. This finding is interesting because NELF complex of proteins that are ubiquitously expressed in testis, associates with DSIF complex to cause transcriptional pausing. The NELF complex is counteracted by the P-TEFb kinase complex (*49*) and the complex is also involved in coordinating RNA polymerase II pausing, premature termination, and chromatin remodeling to regulate HIV transcription (*50*).

Additionally, BRD2, a member of the BET Bromodomain, along with AKAP9, ARHGEF28, and INTS1 are amongst the proteins identified as part of the interactomes of Tat and NF-κB p65 in SEVs. These set of proteins were linked with over 80 diseases and functions, including decompaction of heterochromatin and inflammatory response of bone marrow-derived macrophages. BRD2 also binds BRD3, BRD4, HIST1H3A, CDC5L and other proteins in a network, with biological processes that include chromatin organization; chromatin remodeling; neural tube closure; nucleosome assembly; protein phosphorylation; regulation of gene expression; regulation of transcription from RNA polymerase II promoter; and spermatogenesis.

AKAP9, ARHGEF28, BRD2, and INTS1 are validated by western blot as interactomes of Tat and NF-κB p65 in SEVs. Co-immunoprecipitation and western blot validation assays indicate BRD2, HEXIM 1, and NELFB physically interact with Tat. INTS1 is a member of the integrator complex that regulates and attenuates transcription at promoters that are bound by pausing NELF and DSIF (*51*). The integrator complex consists of 15 family members that perform various functions (cleavage, phosphatase, and core), some of which have been shown to reverse HIV latency (*40, 52*). Given that knockout of INTS12 was shown to increase HIV latency reversal (*53*), our data, showing that NELFB physically interacts with INTS1, is very interesting. Since INTS proteins, such as INTS12 are present in chromatin at the promoter of HIV, it is likely that their effects on HIV may be direct (*53*). That said, it is yet to be determined whether cellular or SEVs-associated INTS1 is present on chromatin at the promoter of HIV and if it has any effect on HIV infection and latency. Future study is required to more fully characterize the interactomes of INTS1 and NELFB in SEVs, determine if other Integrator and/or NELF complexes have physical interactions, as well as determine whether the complexes participate in transcription.

The findings of this current study identify the interactomes of Tat and NF-κB p65 in SEVs and implicate SEVs interactome in HIV transcription. The identification of HEXIM 1, PTBP1, NELFB, INTS1, and BRD2 as interactomes of Tat and NF-κB p65 in SEVs provide potential molecular mechanisms regulating SEVs-mediated HIV transcription and may provide clues on SEVs-mediated chromatin remodeling, transcription initiation and elongation to target Tat-mediated LTR-driven transcription. These findings are significant since permanent reduction of the HIV reservoir has not been achieved, in part, because viral latency involving some aspects of viral transcription are unresponsive to current ART.

## MATERIALS AND METHODS

### Ethical approvals

Specimens collected with the approval of The University of Iowa Institutional Review Board (IRB) were used. HIV-negative and -positive subjects consented to participate in this study via written informed consent. All specimens were received unlinked to any identifiers. All experiments were performed in accordance with the approved University guidelines and regulations.

### Isolation of SEVs

Semen samples were stored at −80°C until isolation. HIV-donors had no history of hepatitis B virus (HBV) and hepatitis C virus (HCV) infection. HIV+ donors were negative for HBV and HCV. SEVs were isolated using a previously described protocol (*54–56*). Briefly, semen samples were thawed at room temperature (RT) and differentially centrifuged at 500× *g* for 10 min, 2000× *g* for 10 min, and 10,000× *g* for 30 min to remove spermatozoa, cellular debris, and large materials, respectively. Supernatants were aliquoted to process further for isolation of SEVs using Particle Purification Liquid Chromatography (PPLC). The 100 cm PPLC Econo-Columns^®^ (Bio-Rad, Hercules, CA, USA) was packed with size-exclusion dextran-based Sephadex™ beads of sizes as follows: G-10 (17-0010-01), G-15 (170020-01), G-25 fine (17-0032-01), G-50 fine (17-0042-01), G-75 (17-0050-01), and G-100 (17-0060-01) purchased from Cytiva (formerly GE Healthcare, Marlborough, MA, USA) using our previously published protocol (*56*). SEVs were isolated at room temperature by gravity. 0.1X phosphate-buffered saline (PBS) was used as the mobile phase. Fractions were collected in Greiner UV-Star^®^ 96-well plates using FC204 fraction collector (Gilson, Middleton, WI, USA), with 6 drops per well. UV-Vis and fluorescence spectral of the fractions were measured using a Synergy H1 plate reader (Biotek, Winooski, VT, USA). Based on 280 nm absorbance, peak 1 and peak 2 (Fraction 1 and 2) were collected and used as SEVs.

### Cells and the Tat plasmid

TZM-bl cells were obtained through the NIH AIDS Reagent Program and were maintained in DMEM (Gibco-BRL/Life Technologies) containing 5% EV-depleted FBS (Gibco), 100 U/ml penicillin, 100 μg/ml streptomycin, sodium pyruvate and 0.3 mg/ml L-glutamine (Invitrogen, Molecular Probes) as previously described (*11*). pCMV Tat plasmid (p-Tat) was a kind gift from Dr. Francesca Di Nunzio (Institut Pasteur, France).

### LTR promoter activation assay

Vehicle (PBS), 100 μg/mL SEVs from HIV-ART-, HIV+ART-, HIV+ART+ were added to TZM-bl indicator cells. Cells were cultured for 24 hours and then assessed for HIV promoter transactivation by measuring luciferase reporter activity using Steady-Glo (Promega) as previously described (*12*).

### SEV-mediated inhibition of Tat

TZM-bl cells were transfected with pTat plasmid using EndoFectin™ Max Transfection Reagent (GeneCopoeia™) at a ratio of 2:1 (µl EndoFectin to µg pTat) in reduced serum media (Opti-MEM™) using our previously described protocol (*12*). Cells were cultured for 3 hours at which point, the medium was replaced with complete DMEM containing 100 μg/mL SEVs or equivalent volume of vehicle (PBS). The effects of SEVs on Tat were determined 24 hours later by measuring promoter activity using the Steady-Glo^®^ (Promega) luminescence assay as previously described (*12*).

### Identification of Tat binding SEV proteins

Bl21 competent *E. coli* were transformed with a plasmid expressing His-Tat, and batches were cultured in 500 mL at 37 °C. In the exponential growth phase, transcription and protein expression were induced with 1 mM Isopropyl β-d-1-thiogalactopyranoside (IPTG) overnight at 16 °C. Cells were lysed with the sonication method and the supernatant was clarified by centrifugation at 10,000 g for 20 minutes. His-tat protein was purified on magnetic DynabeadsTM (ThermoFisher, cat#10103D) to which 200 µg SEVs were added and incubated for 30 minutes rotating at 4 °C. The beads were washed 4 times and proteins were eluted with Laemmli buffer solution at 95 °C for 5 min. A negative control with no *E. coli* extract was performed in parallel. The BLUEstainTM 2 Protein ladder (a three-color protein standard with 13 prestained proteins, covering a wide range molecular weights from 3.5 to 245 kDa) was used as molecular marker. HIV Tat bands (monomers and multimers) (*57, 58*) were cut from the gel and digested with trypsin. Tat-associated SEV proteins were identified by mass spectrometry using the shotgun method (described below).

### Identification of NF-κB binding SEV proteins

10 µg SEV proteins or 100 µg BSA protein were incubated with 1 µM dI:dC (ThermoFisher, cat# 20148E) for cold competitive binding for 20 minutes on ice. After 20 min, 20 picomol NF-kB Cy5-labelled probe was added and the mixture was incubated on ice for another 20 min. The mixture was subjected to 5 % non-denaturing polyacrylamide gel. The shifted band was cut from the gel and eluted in Tris buffer and trypsin-digested. The NF-κB binding proteins were identified by mass spectrometry, described below.

### Western blot analysis

SEVs lysates were separated and processed for western blotting as previously described. The lysates were probed with appropriate primary antibodies. The appropriate secondary IRDye antibodies were used for imaging by Odyssey Infrared Imaging (LI-COR Biosciences) (*54, 59–61*). Image J was used to quantify blots.

### Automated capillary immunoblot analysis

All reagents used in this assay were supplied by ProteinSimple, with the exception of primary antibodies. The ProteinSimple Jess system performs automated size-based separation, detection, and characterization of protein molecular weights in denatured protein lysates using only nanoliter amounts of sample. To quantify the absolute amount of different antigens in SEVs, we followed the manufacturer’s recommended method for 12-230-kDa Jess separation module (SM-W004). Protein extracts (1 μg/μL) in 3 μL from uninfected subjects, HIV+ART-, and HIV+ART+ SEVs were mixed with 0.1X Sample buffer and Fluorescent 5X Master mix (ProteinSimple) to achieve a final concentration of 0.25 μg/μL in the presence of fluorescent molecular weight markers and 400 mM dithiothreitol (ProteinSimple). The mixture was denatured at 95°C for 5 min. Ladder (12-230-kDa PS-ST01EZ) and SEVs proteins were separated in capillaries as they migrated through a separation matrix at 375 volts. A ProteinSimple proprietary photoactivated capture chemistry was used to immobilize separated SEVs proteins on the capillaries. Primary antibodies against AKAP9 (Invitrogen, cat# PA5-113313, polyclonal antibody), ARHGEF28 (Invitrogen, cat# PA558032, polyclonal antibody), BRD2 (Invitrogen, cat# MA534812, recombinant rabbit monoclonal antibody, JG40-42), INTS1 (Antibodies Online, AA 451-710), CD9 (Cell signaling, cat# D8O1A, monoclonal antibody), CD63 (Developmental Studies Hybridoma Bank (DSHB), cat# H5C6, monoclonal antibody), CD81 (Proteintech, cat# 66866-1-Ig, monoclonal antibody), Flotillin (Proteintech, cat# 15571-1-AP, polyclonal antibody), HEXIM1 (D5Y5K) (Cell signaling, Cat#12604, monoclonal antibody), NELFB (D6K9A) (Cell signaling, Cat#14894, monoclonal antibody), were diluted at a 1:100, added to the microplate (SM-FL004) top wells, and incubated for 60 min. After a wash step, anti-rabbit secondary horseradish peroxidase (HRP)-conjugated antibody (042-206) was added for 30 min and chemiluminescent was established with peroxide/luminol-S (ProteinSimple). Apparent molecular weights were calculated through reference to the capillary positions of the ladder peaks from a supplied standard with known molecular weights (**Figs. S1, A and B**). Quantification of signal intensity was calculated using the gaussian method provided in Compass software (version 4.1.0, Protein Simple), which captured the digital images of chemiluminescence of the capillary. Chemiluminescence heights (chemiluminescence intensity), area, and signal/noise ratio were calculated automatically. Results are visualized as electropherograms representing peak of chemiluminescence intensity and as lane view from signal of chemiluminescence detected in the capillary (**Figs. S1, C to L**).

### Mass spectrometry (MS) analysis

MS analysis was performed as described (*55, 56, 62*). Briefly, peptides were resuspended in 5 μL of 0.5% FA and loaded onto a 3-phase MudPIT column (150 μm x 2 cm C18 resin, 150 μm x 4 cm strong cation exchange SCX resin, filter union, and 100 μm x 12 cm C18 resin) as described previously (*63*). A 10-step MudPIT (0 mm, 25 mm, 50 mm, 100 mm, 150 mm, 200 mm, 300 mm, 500 mm, 750 mm, and 1000 mm ammonium acetate, each salt pulse followed by a 120-min acetonitrile gradient 5–50% B [Buffer A: 0.1% FA; Buffer B: 0.1% FA in acetonitrile]) was executed for LC-MS analysis using an Eksigent™ AS-1 autosampler and Eksigent™ nano-LC Ultra 2D pump online with an Orbitrap LTQ XL linear ion trap mass spectrometer (Thermo Finnigan) with a nanospray source. MS data acquisition was done in a data-dependent 6-event method (a survey FTMS scan [res. 30,000] followed by five data-dependent IT scans for five consequent most abundant ions). The general mass spectrometric settings: spray voltage, 2.4 kV; no sheath and no auxiliary gas flow; ion transfer tube temperature, 200 °C; CID fragmentation (for MS/MS), 35% normalized collision energy; activation q = 0.25; activation time, 30 ms. The minimal threshold for the dependent scans was set to 1000 counts, and a dynamic exclusion list was used with the following settings: repeat count of 1, repeat duration of 30 s, exclusion list size of 500, exclusion duration of 90 s.

### Electrophoretic mobility shift assay (EMSA)

EMSA assay was conducted with NF-κB p65 consensus oligonucleotide: forward 5’–Cy5-AGTTGA<u>GGGGACTTTCCC</u>AGGC–3’ and reverse 3’–TCAACTCCCCTGAAAGGGTCCG–5’ with the underlined regions being the binding sites for NF-κB, using our previously described protocol (*12*).

#### Immunoprecipitation assay

Immunoprecipitation assay was carried out as recommended by antibody vendor – Cell signaling and modified per experimental requirements. 250 ug of SEVs used as input was pre-cleared to reduce non-specific protein binding to the Protein G Magnetic beads. Protein G magnetic beads (Cell Signaling Technology, Cat# 70024) were used to pull down NELF using anti-NELFB rabbit mAb (Cell Signaling Technology, Cat #14894). Briefly, 20 μL of Protein G Magnetic beads slurry was prewashed with 1X PBS and then 200 μL of the 250 µg of SEVs was added to 20 μL of pre-washed magnetic beads and incubated with rotation for 20 minutes at room temperature. After 20 min, the beads were separated from the SEVs input using a magnetic separation rack. The pre-cleared SEVs input was transferred to a clean tube and magnetic bead pellet was discarded. 1µg of anti-NELFB antibody was added to 200 μL of pre-cleared SEVs input and incubated with rotation overnight at 4°C to form the immunocomplex. Next morning, SEVs input and antibody (immunocomplex) solution was transferred to the tube containing the pre-washed magnetic bead pellet and incubated with rotation for 20 min at room temperature. After 20 min, the beads were pelleted using a magnetic separation rack. The beads pellet was washed five times with 500 μL of 1X cell lysis buffer. After five washes, the pellet was resuspended with 20 to 40 μL 3X SDS sample buffer. The mixture was vortexed, briefly centrifuged, and heated to 95 to 100°C for 5 min. Beads were pelleted using magnetic separation rack. The supernatant was transferred to a new tube and used for 12% SDS-PAGE gel and Western blot.

### Cell viability

Viability was determined by MTT assay as previously described (*64*).

### Statistics

Two-tailed, paired, student’s t test p-value (GraphPad Prism) calculations determined statistical significance where **** p < 0.0001, *** p < 0.001, ** p <0.01, * p < 0.05, ns = non-significant. Error bars represent standard error of the mean (SEM) across independent experiments unless otherwise noted.

## Supplementary Materials

Figures S1 – S2

Tables S1 – S10

Data files S1 – S2

## Funding

National Institutes of Health grant R01DA042348-01 (to CMO)

National Institutes of Health grant R01DA050169 (to CMO)

National Institutes of Health grant R21/R33DA053643 (to CMO)

VA Merit Review BX000207 (to JTS)

Startup funds from NYMC and Lovelace Biomedical

## Author contributions

Conceptualization: CMO

Methodology: BCO, HK, WN, VP, AC, AN, JTS, CMO

Investigation: HK, WN, VP, AC, AN

Visualization: HK, WN, VP, AC, AN

Funding acquisition: CMO, JTS

Project administration: BCO

Supervision: CMO

Writing – original draft: BCO, HK, WN, VP, AC, AN, JTS, CMO

Writing – review & editing: BCO, HK, WN, VP, AC, AN, JTS, CMO

## Competing interests

Authors declare that they have no competing interests.

## Data and materials availability

The mass spectrometry data are deposited to the ProteomeXchange Consortium via the PRIDE (*65*) partner repository with the dataset identifier PXD062784 and 10.6019/PXD062784 (for NF-κB) and the dataset identifier PXD062816 and 10.6019/PXD062816 (for TAT)” (http://www.ebi.ac.uk/pride). All other data are available in the main text or the supplementary materials.

## Notes

### Competing Interest Statement

The authors have declared no competing interest.

