## Supplemental Material for "Identification of HIV Tat and NF-κB binding proteins associated with semen-derived extracellular vesicles"

**Supplementary Materials**
**Figures S1 – S2**
**Tables S1 – S10**
**Data files S1 – S2**

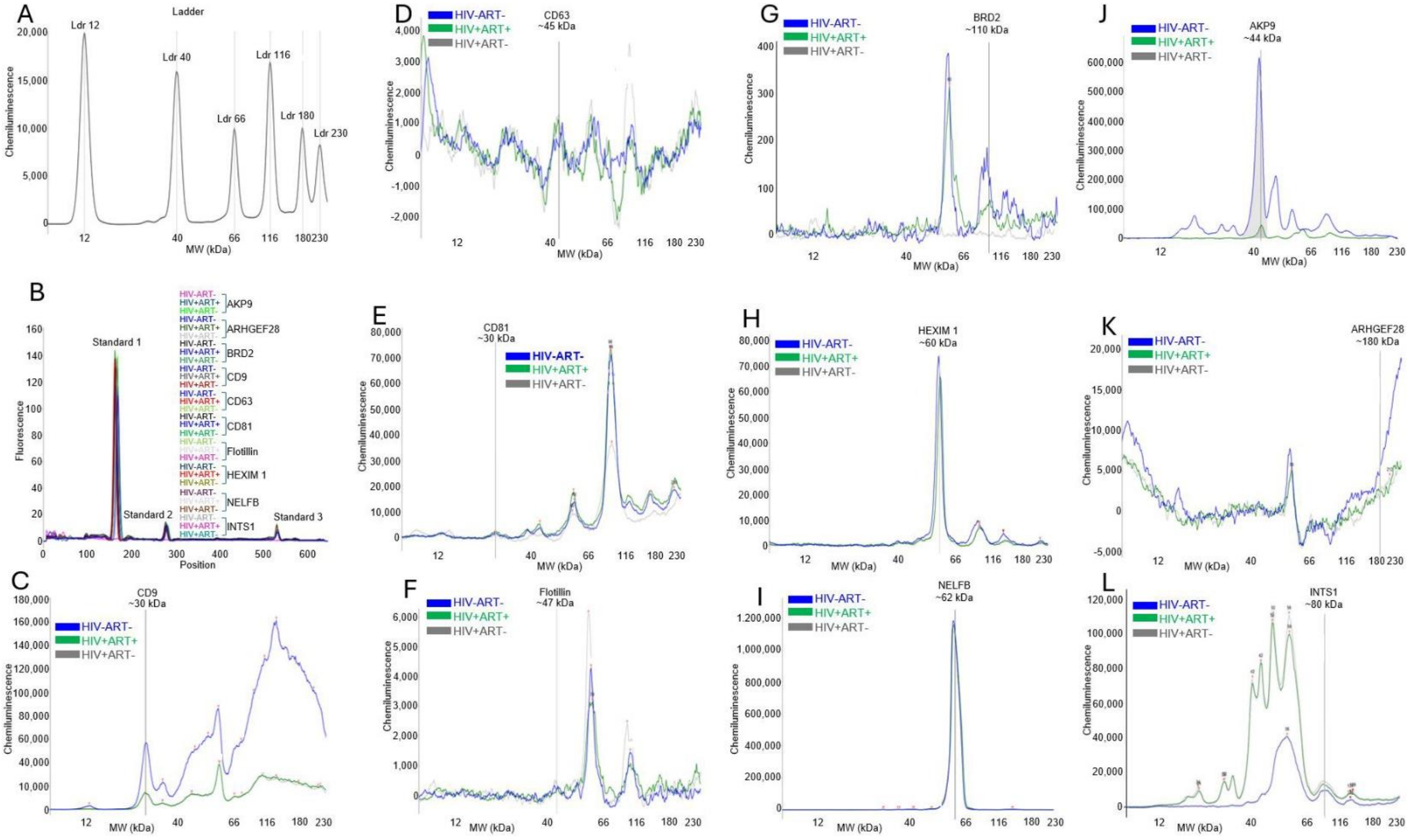

**Figure S1: Capillary-based assay graph visualization of EVs-associated proteins. A**) Electropherograms of biotin ladder. **B**) Fluorescence standard. **C to L**) Electropherograms of SEVs-associated proteins. Area under the peak (marked by a gray vertical line) represents the intensity of the chemiluminescent signal. Images were generated by Jess$^{TM}$ Simple Western system for immunodetection.

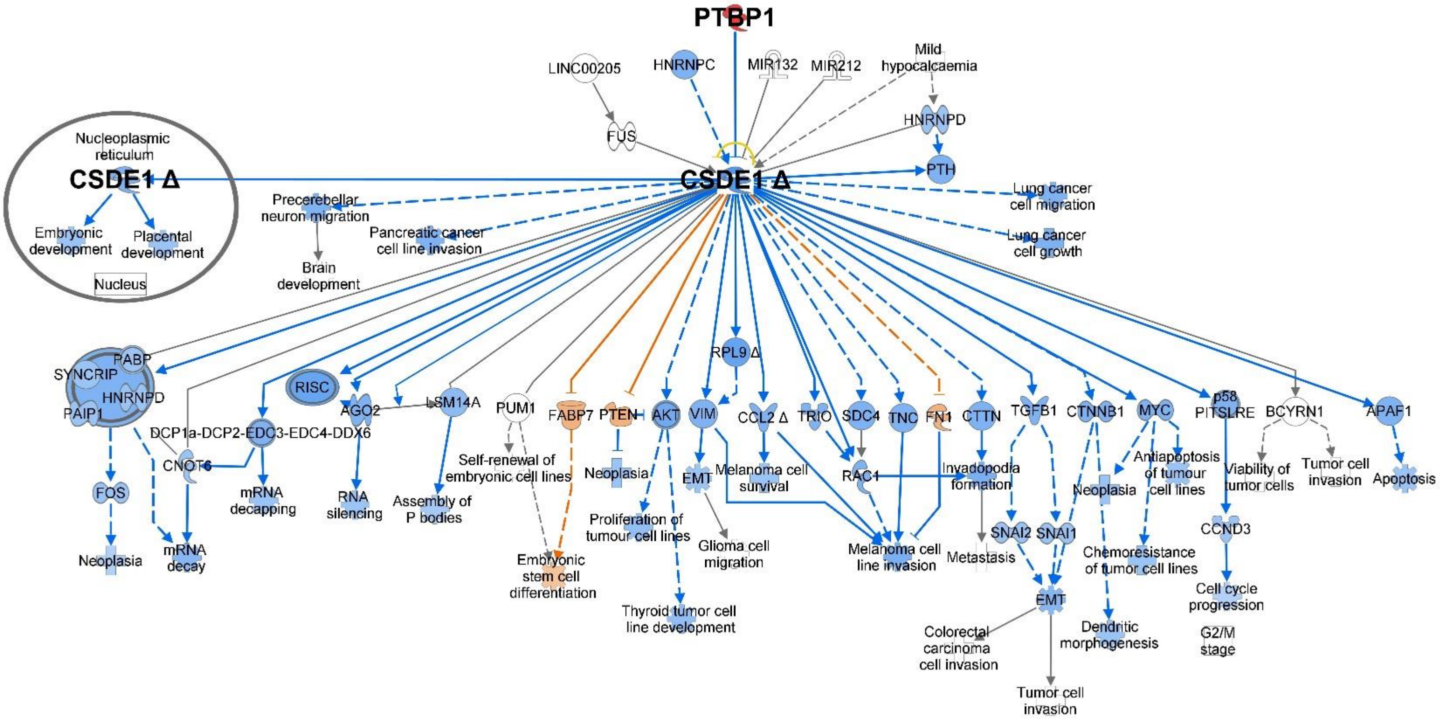

**Figure S2: The PTBP1 interactome.** A model of the interactome of PTBP1 depicting PTBP1-mediated regulation of cold shock domain containing E1 (CSDE1). The pathway was generated with Ingenuety Pathway Analysis (IPA) Path Designer.

| Experiment 1 | | | Experiment 2 | | |
|---|---|---|---|---|---|
| SEV Total (194 proteins) | SEV elution (178 proteins) | Interactomes of Tat in SEVs elution (60 proteins) | SEV Total (183 proteins) | SEV elution (161 proteins) | Interactomes of Tat in SEVs elution (54 proteins) |
| FN1 | FN1 | FN1 | FN1 | FN1 | FN1 |
| LTF | LTF | LTF | LTF | LTF | LTF |
| SEMG2 | SEMG2 | SEMG2 | SEMG2 | SEMG2 | SEMG2 |
| SEMG1 | SEMG1 | SEMG1 | SEMG1 | SEMG1 | SEMG1 |
| TGM4 | TGM4 | TGM4 | TGM4 | TGM4 | TGM4 |
| CLU | CLU | CLU | CLU | CLU | CLU |
| ALB | ALB | ALB | ALB | ALB | ALB |
| MUC6 | MUC6 | MUC6 | MUC6 | MUC6 | MUC6 |
| PIP | PIP | PIP | PIP | PIP | PIP |
| LAMB2 | LAMB2 | KLK3 | LAMB2 | ACPP | FASN |
| ACPP | ACPP | FASN | ACPP | SORD | DPP4 |
| ANPEP | SORD | LPL | ANPEP | KLK3 | CAMP |
| SORD | KLK3 | DPP4 | SORD | PLOD1 | LAMA5 |
| MME | PLOD1 | HSPA8 | MME | LGALS3BP | AKR1B1 |
| KLK3 | LGALS3BP | ANXA2 | KLK3 | ACTB | DCXR |
| LGALS3BP | ACTB | PGC | LGALS3BP | ACTG1 | QSOX1 |
| ACTB | ACTG1 | ACE | ACTB | FASN | ACE |
| ACTG1 | FASN | EDDM3A | ACTG1 | CKB | CLTC |
| FASN | CKB | PYGB | FASN | PLA1A | H2AFX |
| CKB | PLA1A | PFKL | CKB | HSP90AA1 | HIST1H2AA |
| PLA1A | HSP90AA1 | PPP1R7 | PLA1A | LPL | PYGB |
| HSP90AA1 | LPL | SERPINB6 | HSP90AA1 | CPZ | SYNE1 |
| LPL | CPZ | SYNE1 | LPL | HSP90AB1 | GPD1L |
| CPZ | HSP90AB1 | GPD1L | CPZ | SERPINA5 | ANXA11 |
| HSP90AB1 | SERPINA5 | ANXA11 | HSP90AB1 | DPP4 | PYGL |
| SERPINA5 | DPP4 | ACTN1 | SERPINA5 | RAB3D | AKAP9 |
| DPP4 | RAB3D | PYGL | DPP4 | GSTM3 | NAGLU |
| RAB3D | IGHG1 | AKAP9 | RAB3D | HSPA1A | SPG11 |
| IGHG1 | GSTM3 | SPG11 | IGHG1 | HSPA1B | ANKHD1 |
| GSTM3 | HSPA1A | ANK3 | GSTM3 | RAB3B | ANK3 |
| AZGP1 | HSPA1B | SPTBN5 | AZGP1 | CAMP | DCHS1 |
| HSPA1A | RAB3B | EVPL | HSPA1A | ACTA1 | EVPL |
| HSPA1B | CAMP | USP24 | HSPA1B | ACTC1 | DCHS2 |
| RAB3B | ACTA1 | PRR14L | RAB3B | EEF1A1 | GVINP1 |
| CAMP | ACTC1 | GVINP1 | CAMP | EEF1A1P5 | ARHGEF11 |
| ACTA1 | EEF1A1 | ARHGEF11 | ACTA1 | PKM | GPR98 |
| ACTC1 | EEF1A1P5 | TTC21A | ACTC1 | LAMA5 | TTC21A |
| EEF1A1 | PKM | LRRIQ1 | EEF1A1 | AKR1B1 | LRRIQ1 |
| EEF1A1P5 | LAMA5 | TBC1D15 | EEF1A1P5 | HSPA8 | CDH19 |
| PKM | AKR1B1 | FRMD6-AS1 | PKM | TUBA1C | PCNX3 |
| IGHA1 | HSPA8 | PFAS | IGHA1 | TUBA1A | GPR34 |
| LAMA5 | ANXA1 | FAT3 | LAMA5 | SIL1 | FAT3 |
| LAMC1 | TUBA1C | ADGRA2 | AKR1B1 | CRISP1 | PCDH1 |
| PDCD6IP | TUBA1A | ARHGEF28 | LAMC1 | TUBA1B | WDR87 |
| HSPA8 | SIL1 | PTPRQ | PDCD6IP | GAPDH | ARHGEF28 |
| IDH1 | MATN2 | GNPTAB | HSPA8 | RAB27A | RYR3 |
| ANXA1 | IGKC | IL7R | IDH1 | TIMP1 | PTPRQ |
| TUBA1C | CRISP1 | ITSN1 | ANXA1 | SCPEP1 | GNPTAB |
| TUBA1A | TUBA1B | MAPKBP1 | TUBA1C | PRDX6 | ITSN1 |
| SIL1 | GAPDH | TRIM33 | TUBA1A | DCXR | PTBP1 |
| MATN2 | RAB27A | PTBP1 | SIL1 | PLOD3 | INTS1 |
| IGKC | TIMP1 | NAALAD2 | MATN2 | QSOX1 | GSTCD |
| CRISP1 | SCPEP1 | INTS1 | IGKC | GSTP1 | CCDC157 |
| TUBA1B | PRDX6 | MARS2 | CRISP1 | LIPG | BRD2 |
| GAPDH | DCXR | GSTCD | TUBA1B | MAMDC2 | |
| RAB27A | PLOD3 | DNAH5 | GAPDH | YWHAZ | |
| GDI2 | QSOX1 | CCDC157 | RAB27A | MSMB | |
| ANXA2 | GSTP1 | UGT1A7 | GDI2 | PGC | |
| TIMP1 | LIPG | UGT1A8 | ANXA2 | CNDP2 | |
| CTSB | MAMDC2 | BRD2 | TIMP1 | CHID1 | |
| PRDX6 | YWHAZ | | CTSB | PAEP | |
| DCXR | MSMB | | SCPEP1 | VTN | |
| IGHG2 | CD177 | | PRDX6 | PRCP | |
| QSOX1 | PGC | | DCXR | HIST1H2BA | |
| GSTP1 | CNDP2 | | IGHG2 | UBA52 | |
| YWHAZ | CHID1 | | QSOX1 | RPS27A | |
| MSMB | PAEP | | GSTP1 | UBB | |
| CD177 | CST4 | | YWHAZ | UBC | |

| | | | | |
|---|---|---|---|---|
| PGC | VTN | | MSMB | NPNT |
| CNDP2 | PRCP | | CD177 | ACE |
| PAEP | HIST1H2BA | | PGC | LGMN |
| CST4 | UBA52 | | CNDP2 | FAM129A |
| HIST1H2BA | RPS27A | | CHID1 | CLTC |
| UBA52 | UBB | | PAEP | CCT5 |
| RPS27A | UBC | | CST4 | GLA |
| UBB | NPNT | | HIST1H2BA | YWHAG |
| UBC | ACE | | UBA52 | EDDM3A |
| ACE | LGMN | | RPS27A | HSP90B1 |
| FAM129A | FAM129A | | UBB | PATE1 |
| ANXA5 | CLTC | | UBC | GDF15 |
| EZR | CCT5 | | FAM129A | CCT4 |
| YWHAG | ANXA5 | | ANXA5 | H2AFX |
| EDDM3A | GLA | | EZR | HIST1H2AA |
| HSP90B1 | EZR | | YWHAG | PRDX4 |
| IGLC3 | EDDM3A | | EDDM3A | CAND1 |
| IGLC2 | HSP90B1 | | HSP90B1 | SMPD1 |
| IGLC6 | PATE1 | | PATE1 | YWHAE |
| GDF15 | GDF15 | | IGLC3 | PRDX2 |
| PPIA | CCT4 | | IGLC2 | HIST1H2BL |
| TUBB4B | H2AFX | | IGLC6 | HIST1H2BM |
| PRDX4 | HIST1H2AA | | GDF15 | HIST1H2BK |
| YWHAE | PRDX4 | | PPIA | HIST2H2BE |
| PRDX2 | CAND1 | | TUBB4B | HIST1H2BH |
| HIST1H2BL | SMPD1 | | YWHAE | HIST1H2BB |
| HIST1H2BM | YWHAE | | PRDX2 | HIST1H2BC |
| HIST1H2BK | PRDX2 | | HIST1H2BL | HIST1H2BJ |
| HIST2H2BE | HIST1H2BL | | HIST1H2BM | HIST1H2BD |
| HIST1H2BH | HIST1H2BM | | HIST1H2BK | HIST1H2BO |
| HIST1H2BB | HIST1H2BK | | HIST2H2BE | HIST2H2BF |
| HIST1H2BC | HIST2H2BE | | HIST1H2BH | HIST1H2BN |
| HIST1H2BJ | HIST1H2BH | | HIST1H2BB | H2BFS |
| HIST1H2BD | HIST1H2BB | | HIST1H2BC | VWA1 |
| HIST1H2BO | HIST1H2BC | | HIST1H2BJ | HEXA |
| HIST2H2BF | HIST1H2BJ | | HIST1H2BD | VCP |
| HIST1H2BN | HIST1H2BD | | HIST1H2BO | CDH1 |
| H2BFS | HIST1H2BO | | HIST2H2BF | PSAP |
| DOPEY2 | HIST2H2BF | | HIST1H2BN | NME3 |
| PGK1 | HIST1H2BN | | H2BFS | TPP1 |
| VCP | H2BFS | | DOPEY2 | PYGB |
| ENO1 | DOPEY2 | | VWA1 | KLK2 |
| CDH1 | VWA1 | | PGK1 | HIST1H4A |
| PSAP | HEXA | | HEXA | GLB1 |
| CD9 | VCP | | VCP | TF |
| FOLH1 | CDH1 | | PSAP | LDHC |
| PYGB | NME3 | | CD9 | PGD |
| KLK2 | TPP1 | | TPP1 | GANAB |
| HIST1H4A | PYGB | | FOLH1 | DEFA3 |
| TF | PFKL | | PYGB | DEFA1 |
| ANXA3 | KLK2 | | KLK2 | ELSPBP1 |
| AHCY | PPP1CB | | HIST1H4A | CLIC1 |
| GAS6 | HIST1H4A | | TF | PPP1R7 |
| ALDOA | GLB1 | | ANXA3 | GC |
| CPE | TF | | LDHC | MXRA5 |
| DEFA3 | LDHC | | AHCY | PPIB |
| DEFA1 | PGD | | GAS6 | VARS |
| ELSPBP1 | GANAB | | ALDOA | EDDM3B |
| CLIC1 | DEFA3 | | CPE | RAB27B |
| RNASE4 | DEFA1 | | DEFA3 | FUT5 |
| OLFM4 | ELSPBP1 | | DEFA1 | FUT6 |
| LCP1 | CLIC1 | | ELSPBP1 | FUT3 |
| GC | RNASE4 | | CLIC1 | LSR |
| RHOA | GC | | PPP1R7 | RAB15 |
| CST3 | EEF2 | | OLFM4 | CTSH |
| SYTL1 | PPIB | | LCP1 | LCN15 |
| EEF2 | VARS | | GC | YWHAB |
| ALOX15B | EDDM3B | | RHOA | MANBA |
| RAB27B | RAB27B | | CST3 | GAPDHS |
| PEBP1 | CNP | | SYTL1 | CCT2 |
| B2M | LSR | | EDDM3B | C1QTNF1 |
| RHOC | RAB15 | | RAB27B | SYNE1 |
| ZG16B | LCN15 | | PEBP1 | ADAMTS1 |
| CNP | RAB2A | | ZG16B | PATE4 |

| | | | | |
|---|---|---|---|---|
| LSR | MANBA | | CNP | YWHAQ |
| RAB15 | ASAH1 | | RAB15 | CA2 |
| GLIPR2 | GAPDHS | | GLIPR2 | ANXA11 |
| PLPP1 | CCT2 | | PLPP1 | PRDX1 |
| TMPRSS2 | C1QTNF1 | | SERPINB6 | EPHX2 |
| CD63 | SYNE1 | | PIGR | CES5A |
| PGAM1 | ADAMTS1 | | TMPRSS2 | NAGLU |
| FBLN2 | PATE4 | | CD63 | CPQ |
| YWHAB | YWHAQ | | YWHAB | CELSR3 |
| GLG1 | PSMA2 | | ASAH1 | ACR |
| ASAH1 | MMP2 | | SYNE1 | USP24 |
| SYNE1 | GLB1L | | NPC2 | FUCA1 |
| NPC2 | CA2 | | YWHAQ | ARHGEF11 |
| GPD1L | ANXA11 | | ANXA11 | TTC21A |
| YWHAQ | EPHX2 | | PRDX1 | OS9 |
| PSMA2 | ZPBP | | MPO | ANKRD18B |
| PRSS8 | PYGL | | FCGBP | PARP3 |
| ANXA11 | CTSD | | EPHX2 | FAT3 |
| PRDX1 | CES5A | | PFN1 | TRPM6 |
| MPO | NAGLU | | CTSD | |
| EPHX2 | ECM1 | | PSMA1 | |
| CFL1 | CPQ | | NAGLU | |
| TMBIM1 | CELSR3 | | ECM1 | |
| NAGLU | ACR | | ANK3 | |
| GNAS | DCHS2 | | SLPI | |
| GNAS | FUCA1 | | ARF3 | |
| ECM1 | SEMA3F | | ARF1 | |
| CMPK1 | ARHGEF11 | | DCHS1 | |
| GPI | TTC21A | | PRKAR2A | |
| ANK3 | PGK2 | | USP24 | |
| SLPI | ANKRD18B | | TSPAN1 | |
| ARF3 | FAT3 | | COL6A1 | |
| ARF1 | KIF5B | | ARHGEF11 | |
| PSMA6 | TRPM6 | | TTC21A | |
| CELSR3 | HSPA13 | | ACTR3 | |
| RAC2 | BDP1 | | ANKRD18B | |
| RAC1 | | | FAT3 | |
| PRKAR2A | | | MAP1B | |
| COL6A1 | | | TRPM6 | |
| ARHGEF11 | | | CAPN1 | |
| TTC21A | | | CAPZB | |
| C3 | | | | |
| FRMD6-AS1 | | | | |
| CDC42 | | | | |
| FBP1 | | | | |
| ANKRD18B | | | | |
| GPR34 | | | | |
| FAT3 | | | | |
| MAP1B | | | | |
| TRPM6 | | | | |
| SCN11A | | | | |
| INTS6L | | | | |

**Table S1: Interactomes of Tat in SEVs.** The proteins were identified using mass spectrometric (MS) analysis. Data were generated from 2 independent experiments.

| Elements in 2-way Venn for Experiment 1 | | | Elements in 2-way Venn for Experiment 2 | | | Elements in 2-way Venn for combined Experiments 1 and 2 | | |
|---|---|---|---|---|---|---|---|---|
| **153 elements included exclusively in "SE Elution 178":** | **25 common elements in "S2 178" and "S3 60":** | **35 elements included exclusively in "Tat\|SE Elution60":** | **134 elements included exclusively in "SE Elution 161":** | **27 common elements in "S5 161" and "S6 54":** | **27 elements included exclusively in "Tat\|SE Elution54":** | **16 common elements in "Tat\|SE Elution35" and "Tat\|SE Elution27":** | **19 elements included exclusively in "Tat\|SE Elution35":** | **11 elements included exclusively in "Tat\|SE Elution27":** |
| LAMB2 | FN1 | ANXA2 | ACPP | FN1 | GPD1L | EVPL | ANXA2 | PYGL |
| ACPP | LTF | PPP1R7 | SORD | LTF | PYGL | GVIN1 | PPP1R7 | ANKHD1 |
| SORD | SEMG2 | SERPINB6 | KLK3 | SEMG2 | AKAP9 | PTBP1 | SERPINB6 | DCHS1 |
| PLOD1 | SEMG1 | GPD1L | PLOD1 | SEMG1 | SPG11 | AKAP9 | ACTN1 | DCHS2 |
| LGALS3BP | TGM4 | ACTN1 | LGALS3BP | TGM4 | ANKHD1 | SPTCS | SPTBN5 | GPR98 |
| ACTB | CLU | AKAP9 | ACTB | CLU | ANK3 | ARG28 | USP24 | CDH19 |
| ACTG1 | ALB | SPG11 | ACTG1 | ALB | DCHS1 | PTPRQ | PRR14L | PCNX3 |
| CKB | MUC6 | ANK3 | CKB | MUC6 | EVPL | GSTCD | TBC1D15 | GPR34 |
| PLA1A | PIP | SPTBN5 | PLA1A | PIP | DCHS2 | ITSN1 | FRMD6-AS1 | PCDH1 |
| HSP90AA1 | KLK3 | EVPL | HSP90AA1 | FASN | GVINP1 | INT1 | PFAS | WDR87 |
| CPZ | FASN | USP24 | LPL | DPP4 | GPR98 | LRIQ1 | ADGRA2 | RYR3 |
| HSP90AB1 | LPL | PRR14L | CPZ | CAMP | LRRIQ1 | CC157 | IL7R | |
| SERPINA5 | DPP4 | GVINP1 | HSP90AB1 | LAMA5 | CDH19 | GNPTA | MAPKBP1 | |
| RAB3D | HSPA8 | LRRIQ1 | SERPINA5 | AKR1B1 | PCNX3 | GPD1L | TRIM33 | |
| IGHG1 | PGC | TBC1D15 | RAB3D | DCXR | GPR34 | ANK3 | NAALAD2 | |
| GSTM3 | ACE | FRMD6-AS1 | GSTM3 | QSOX1 | PCDH1 | BRD2 | MARS2 | |
| HSPA1A | EDDM3A | PFAS | HSPA1A | ACE | WDR87 | | DNAH5 | |
| HSPA1B | PYGB | ADGRA2 | HSPA1B | CLTC | ARHGEF28 | | UGT1A7 | |
| RAB3B | PFKL | ARHGEF28 | RAB3B | H2AFX | RYR3 | | UGT1A8 | |
| CAMP | SYNE1 | PTPRQ | ACTA1 | HIST1H2AA | PTPRQ | | | |
| ACTA1 | ANXA11 | GNPTAB | ACTC1 | PYGB | GNPTAB | | | |
| ACTC1 | PYGL | IL7R | EEF1A1 | SYNE1 | ITSN1 | | | |
| EEF1A1 | ARHGEF11 | ITSN1 | EEF1A1P5 | ANXA11 | PTBP1 | | | |
| EEF1A1P5 | TTC21A | MAPKBP1 | PKM | NAGLU | INTS1 | | | |
| PKM | FAT3 | TRIM33 | HSPA8 | ARHGEF11 | GSTCD | | | |
| LAMA5 | | PTBP1 | TUBA1C | TTC21A | CCDC157 | | | |
| AKR1B1 | | NAALAD2 | TUBA1A | FAT3 | BRD2 | | | |
| ANXA1 | | INTS1 | SIL1 | | | | | |
| TUBA1C | | MARS2 | CRISP1 | | | | | |
| TUBA1A | | GSTCD | TUBA1B | | | | | |
| SIL1 | | DNAH5 | GAPDH | | | | | |
| MATN2 | | CCDC157 | RAB27A | | | | | |
| IGKC | | UGT1A7 | TIMP1 | | | | | |
| CRISP1 | | UGT1A8 | SCPEP1 | | | | | |
| TUBA1B | | BRD2 | PRDX6 | | | | | |
| GAPDH | | | PLOD3 | | | | | |
| RAB27A | | | GSTP1 | | | | | |
| TIMP1 | | | LIPG | | | | | |
| SCPEP1 | | | MAMDC2 | | | | | |

| | | | |
|---|---|---|---|
| PRDX6 | | | YWHAZ |
| DCXR | | | MSMB |
| PLOD3 | | | PGC |
| QSOX1 | | | CNDP2 |
| GSTP1 | | | CHID1 |
| LIPG | | | PAEP |
| MAMDC2 | | | VTN |
| YWHAZ | | | PRCP |
| MSMB | | | HIST1H2BA |
| CD177 | | | UBA52 |
| CNDP2 | | | RPS27A |
| CHID1 | | | UBB |
| PAEP | | | UBC |
| CST4 | | | NPNT |
| VTN | | | LGMN |
| PRCP | | | FAM129A |
| HIST1H2BA | | | CCT5 |
| UBA52 | | | GLA |
| RPS27A | | | YWHAG |
| UBB | | | EDDM3A |
| UBC | | | HSP90B1 |
| NPNT | | | PATE1 |
| LGMN | | | GDF15 |
| FAM129A | | | CCT4 |
| CLTC | | | PRDX4 |
| CCT5 | | | CAND1 |
| ANXA5 | | | SMPD1 |
| GLA | | | YWHAE |
| EZR | | | PRDX2 |
| HSP90B1 | | | HIST1H2BL |
| PATE1 | | | HIST1H2BM |
| GDF15 | | | HIST1H2BK |
| CCT4 | | | HIST2H2BE |
| H2AFX | | | HIST1H2BH |
| HIST1H2AA | | | HIST1H2BB |
| PRDX4 | | | HIST1H2BC |
| CAND1 | | | HIST1H2BJ |
| SMPD1 | | | HIST1H2BD |
| YWHAE | | | HIST1H2BO |
| PRDX2 | | | HIST2H2BF |
| HIST1H2BL | | | HIST1H2BN |
| HIST1H2BM | | | H2BFS |
| HIST1H2BK | | | VWA1 |
| HIST2H2BE | | | HEXA |
| HIST1H2BH | | | VCP |
| HIST1H2BB | | | CDH1 |

| | | | |
|---|---|---|---|
| HIST1H2BC | | | PSAP |
| HIST1H2BJ | | | NME3 |
| HIST1H2BD | | | TPP1 |
| HIST1H2BO | | | KLK2 |
| HIST2H2BF | | | HIST1H4A |
| HIST1H2BN | | | GLB1 |
| H2BFS | | | TF |
| DOPEY2 | | | LDHC |
| VWA1 | | | PGD |
| HEXA | | | GANAB |
| VCP | | | DEFA3 |
| CDH1 | | | DEFA1 |
| NME3 | | | ELSPBP1 |
| TPP1 | | | CLIC1 |
| KLK2 | | | PPP1R7 |
| PPP1CB | | | GC |
| HIST1H4A | | | MXRA5 |
| GLB1 | | | PPIB |
| TF | | | VARS |
| LDHC | | | EDDM3B |
| PGD | | | RAB27B |
| GANAB | | | FUT5 |
| DEFA3 | | | FUT6 |
| DEFA1 | | | FUT3 |
| ELSPBP1 | | | LSR |
| CLIC1 | | | RAB15 |
| RNASE4 | | | CTSH |
| GC | | | LCN15 |
| EEF2 | | | YWHAB |
| PPIB | | | MANBA |
| VARS | | | GAPDHS |
| EDDM3B | | | CCT2 |
| RAB27B | | | C1QTNF1 |
| CNP | | | ADAMTS1 |
| LSR | | | PATE4 |
| RAB15 | | | YWHAQ |
| LCN15 | | | CA2 |
| RAB2A | | | PRDX1 |
| MANBA | | | EPHX2 |
| ASAH1 | | | CES5A |
| GAPDHS | | | CPQ |
| CCT2 | | | CELSR3 |
| C1QTNF1 | | | ACR |
| ADAMTS1 | | | USP24 |
| PATE4 | | | FUCA1 |
| YWHAQ | | | OS9 |

| | | | |
|---|---|---|---|
| PSMA2 | | | ANKRD18B |
| MMP2 | | | PARP3 |
| GLB1L | | | TRPM6 |
| CA2 | | | |
| EPHX2 | | | |
| ZPBP | | | |
| CTSD | | | |
| CES5A | | | |
| NAGLU | | | |
| ECM1 | | | |
| CPQ | | | |
| CELSR3 | | | |
| ACR | | | |
| DCHS2 | | | |
| FUCA1 | | | |
| SEMA3F | | | |
| PGK2 | | | |
| ANKRD18B | | | |
| KIF5B | | | |
| TRPM6 | | | |
| HSPA13 | | | |
| BDP1 | | | |

**Table S2: Elements in the two-way Venn diagram analyses of SEVs proteins Tat interactome.** The elements were identified using freely available, web accessible 2-way Venn diagram. The columns represent proteins unique or common between experiments. Data were generated from 2 independent experiments. Blue represents proteins common between the two experiments.

| From Molecule | Relationship Type | To Molecule(s) |
| --- | --- | --- |
| **AKAP9** | localization | IL2 |
| **AKAP9** | protein-protein interactions | AKAP9 |
| Akt | activation | Akt |
| Akt | activation | CTNNB1 |
| Akt | activation | HIF1A |
| Akt | activation | STAT3 |
| Akt | causation | CSDE1 Signaling Pathway |
| Akt | chemical-protein interactions | phosphatidylinositol-3,4,5-triphosphate |
| Akt | expression | Akt |
| Akt | expression | CTNNB1 |
| Akt | expression | GNPTAB |
| Akt | expression | HIF1A |
| Akt | expression | MYC |
| Akt | inhibition | Akt |
| Akt | localization | IL2 |
| Akt | molecular cleavage | CTNNB1 |
| Akt | phosphorylation | Akt |
| Akt | phosphorylation | CTNNB1 |
| Akt | phosphorylation | STAT3 |
| Akt | protein-protein interactions | Akt |
| Akt | protein-protein interactions | CTNNB1 |
| Akt | regulation of binding | CTNNB1 |
| Akt | regulation of binding | HIF1A |
| Akt | regulation of binding | STAT3 |
| Akt | regulation of binding | phosphatidylinositol-3,4,5-triphosphate |
| Akt | translocation | Akt |
| BRD2 | molecular cleavage | BRD2 |
| BRD2 | protein-DNA interactions | MYC |
| BRD2 | protein-protein interactions | BRD2 |
| BRD2 | protein-protein interactions | MYC |
| BRD2 | regulation of binding | BRD2 |
| BRD2 | ubiquitination | BRD2 |
| CTNNB1 | activation | Akt |
| CTNNB1 | activation | CTNNB1 |
| CTNNB1 | activation | MYC |
| CTNNB1 | causation | CSDE1 Signaling Pathway |
| CTNNB1 | expression | CTNNB1 |
| CTNNB1 | expression | GATA3 |
| CTNNB1 | expression | HIF1A |
| CTNNB1 | expression | IL2 |
| CTNNB1 | expression | ITSN1 |
| CTNNB1 | expression | MYC |
| CTNNB1 | expression | STAT3 |
| CTNNB1 | expression | WNT1 |
| CTNNB1 | expression | alanine |
| CTNNB1 | localization | CTNNB1 |
| CTNNB1 | localization | IL2 |
| CTNNB1 | molecular cleavage | CTNNB1 |
| CTNNB1 | phosphorylation | Akt |
| CTNNB1 | phosphorylation | CTNNB1 |
| CTNNB1 | protein-DNA interactions | GATA3 |
| CTNNB1 | protein-DNA interactions | IL2 |
| CTNNB1 | protein-DNA interactions | MYC |
| CTNNB1 | protein-DNA interactions | STAT3 |

| | | |
|---|---|---|
| CTNNB1 | protein-protein interactions | ANK3 |
| CTNNB1 | protein-protein interactions | Akt |
| CTNNB1 | protein-protein interactions | CTNNB1 |
| CTNNB1 | protein-protein interactions | HIF1A |
| CTNNB1 | protein-protein interactions | MYC |
| CTNNB1 | protein-protein interactions | STAT3 |
| CTNNB1 | protein-protein interactions | WNT1 |
| CTNNB1 | regulation of binding | CTNNB1 |
| CTNNB1 | regulation of binding | GATA3 |
| CTNNB1 | regulation of binding | MYC |
| CTNNB1 | regulation of binding | STAT3 |
| CTNNB1 | transcription | MYC |
| CTNNB1 | translocation | CTNNB1 |
| CTNNB1 | ubiquitination | CTNNB1 |
| GATA3 | activation | Akt |
| GATA3 | expression | EVPL |
| GATA3 | expression | GATA3 |
| GATA3 | expression | HIF1A |
| GATA3 | expression | IL2 |
| GATA3 | molecular cleavage | HIF1A |
| GATA3 | phosphorylation | Akt |
| GATA3 | protein-DNA interactions | GATA3 |
| GATA3 | protein-DNA interactions | MYC |
| GATA3 | protein-protein interactions | HIF1A |
| GATA3 | transcription | GATA3 |
| GATA3 | ubiquitination | HIF1A |
| GATA3-AS1 | expression | CTNNB1 |
| GATA3-AS1 | expression | WNT1 |
| GATA3-AS1 | ubiquitination | GATA3 |
| GNPTAB | protein-protein interactions | GNPTAB |
| GPD1L | activation | HIF1A |
| GPD1L | expression | HIF1A |
| GPD1L | modification | HIF1A |
| HIF1A | activation | Akt |
| HIF1A | activation | HIF1A |
| HIF1A | activation | MYC |
| HIF1A | activation | STAT3 |
| HIF1A | expression | HIF1A |
| HIF1A | expression | MYC |
| HIF1A | expression | STAT3 |
| HIF1A | inhibition | CTNNB1 |
| HIF1A | inhibition | HIF1A |
| HIF1A | localization | CTNNB1 |
| HIF1A | localization | HIF1A |
| HIF1A | modification | CTNNB1 |
| HIF1A | modification | HIF1A |
| HIF1A | molecular cleavage | HIF1A |
| HIF1A | molecular cleavage | MYC |
| HIF1A | phosphorylation | Akt |
| HIF1A | phosphorylation | HIF1A |
| HIF1A | phosphorylation | STAT3 |
| HIF1A | protein-DNA interactions | HIF1A |
| HIF1A | protein-DNA interactions | WNT1 |
| HIF1A | protein-protein interactions | CTNNB1 |
| HIF1A | protein-protein interactions | GATA3 |
| HIF1A | protein-protein interactions | HIF1A |
| HIF1A | protein-protein interactions | MYC |

| | | |
|---|---|---|
| HIF1A | protein-protein interactions | STAT3 |
| HIF1A | regulation of binding | CTNNB1 |
| HIF1A | regulation of binding | HIF1A |
| HIF1A | regulation of binding | MYC |
| HIF1A | regulation of binding | STAT3 |
| HIF1A | transcription | HIF1A |
| HIF1A | translocation | HIF1A |
| HIF1A | ubiquitination | HIF1A |
| HOXA11-AS | expression | CTNNB1 |
| HOXA11-AS | expression | STAT3 |
| HOXA11-AS | regulation of binding | CTNNB1 |
| Hmga2 | expression | IL2 |
| IL2 | activation | Akt |
| IL2 | activation | GATA3 |
| IL2 | activation | IL2 |
| IL2 | activation | STAT3 |
| IL2 | expression | GATA3 |
| IL2 | expression | GLS2 |
| IL2 | expression | IL2 |
| IL2 | expression | MYC |
| IL2 | expression | STAT3 |
| IL2 | localization | IL2 |
| IL2 | phosphorylation | Akt |
| IL2 | phosphorylation | GATA3 |
| IL2 | phosphorylation | STAT3 |
| IL2 | protein-protein interactions | IL2 |
| IL2 | regulation of binding | CTNNB1 |
| IL2 | regulation of binding | IL2 |
| IL2 | regulation of binding | STAT3 |
| IL2 | transcription | IL2 |
| IL2 | transcription | MYC |
| IL2 | translocation | IL2 |
| ITSN1 | activation | Akt |
| ITSN1 | protein-protein interactions | ITSN1 |
| Interferon alpha | activation | Akt |
| Interferon alpha | activation | STAT3 |
| Interferon alpha | expression | GATA3 |
| Interferon alpha | expression | GVINP1 |
| Interferon alpha | expression | IL2 |
| Interferon alpha | expression | Interferon alpha |
| Interferon alpha | expression | MYC |
| Interferon alpha | expression | PTBP1 |
| Interferon alpha | expression | STAT3 |
| Interferon alpha | inhibition | Akt |
| Interferon alpha | localization | STAT3 |
| Interferon alpha | phosphorylation | Akt |
| Interferon alpha | phosphorylation | STAT3 |
| Interferon alpha | regulation of binding | MYC |
| Interferon alpha | regulation of binding | STAT3 |
| Interferon alpha | transcription | Interferon alpha |
| Interferon alpha | transcription | STAT3 |
| Interferon alpha | translocation | Interferon alpha |
| Interferon alpha | translocation | STAT3 |
| MALAT1 | activation | Akt |
| MALAT1 | expression | Interferon alpha |
| MALAT1 | phosphorylation | Akt |
| MYC | activation | Akt |

| | | |
|---|---|---|
| MYC | activation | CTNNB1 |
| MYC | activation | MYC |
| MYC | activation | STAT3 |
| MYC | causation | CSDE1 Signaling Pathway |
| MYC | expression | Akt |
| MYC | expression | BRD2 |
| MYC | expression | CTNNB1 |
| MYC | expression | EVPL |
| MYC | expression | GLS2 |
| MYC | expression | HIF1A |
| MYC | expression | MYC |
| MYC | expression | PTBP1 |
| MYC | expression | STAT3 |
| MYC | inhibition | MYC |
| MYC | localization | HIF1A |
| MYC | localization | MYC |
| MYC | molecular cleavage | MYC |
| MYC | phosphorylation | Akt |
| MYC | phosphorylation | MYC |
| MYC | phosphorylation | STAT3 |
| MYC | protein-DNA interactions | MYC |
| MYC | protein-RNA interactions | PTBP1 |
| MYC | protein-protein interactions | BRD2 |
| MYC | protein-protein interactions | CTNNB1 |
| MYC | protein-protein interactions | DNTT |
| MYC | protein-protein interactions | HIF1A |
| MYC | protein-protein interactions | MYC |
| MYC | protein-protein interactions | PTBP1 |
| MYC | regulation of binding | Akt |
| MYC | regulation of binding | CTNNB1 |
| MYC | regulation of binding | HIF1A |
| MYC | regulation of binding | MYC |
| MYC | transcription | CTNNB1 |
| MYC | transcription | DNTT |
| MYC | transcription | MYC |
| MYC | ubiquitination | MYC |
| NOVA2 | protein-protein interactions | BRD2 |
| NOVA2 | protein-protein interactions | NOVA2 |
| PTBP1 | causation | CSDE1 Signaling Pathway |
| PTBP1 | expression | CTNNB1 |
| PTBP1 | expression | GLS2 |
| PTBP1 | expression | IL2 |
| PTBP1 | expression | MYC |
| PTBP1 | expression | alanine |
| PTBP1 | protein-DNA interactions | Hmga2 |
| PTBP1 | protein-RNA interactions | CTNNB1 |
| PTBP1 | protein-RNA interactions | HIF1A |
| PTBP1 | protein-RNA interactions | HOXA11-AS |
| PTBP1 | protein-RNA interactions | MALAT1 |
| PTBP1 | protein-RNA interactions | MYC |
| PTBP1 | protein-protein interactions | DNTT |
| PTBP1 | protein-protein interactions | MYC |
| PTBP1 | protein-protein interactions | NOVA2 |
| PTBP1 | protein-protein interactions | PTBP1 |
| PTPRQ | activation | phosphatidylinositol-3,4,5-triphosphate |
| PTPRQ | phosphorylation | phosphatidylinositol-3,4,5-triphosphate |
| PTPRQ | protein-protein interactions | CTNNB1 |

| | | |
|---|---|---|
| RAVER2 | molecular cleavage | RAVER2 |
| RAVER2 | protein-protein interactions | GATA3 |
| RAVER2 | protein-protein interactions | PTBP1 |
| RBM24 | protein-protein interactions | EVPL |
| RBM24 | protein-protein interactions | MYC |
| STAT3 | activation | Akt |
| STAT3 | activation | CTNNB1 |
| STAT3 | activation | Interferon alpha |
| STAT3 | activation | MYC |
| STAT3 | activation | STAT3 |
| STAT3 | expression | GATA3 |
| STAT3 | expression | HIF1A |
| STAT3 | expression | IL2 |
| STAT3 | expression | Interferon alpha |
| STAT3 | expression | MYC |
| STAT3 | expression | STAT3 |
| STAT3 | expression | mir-124 |
| STAT3 | inhibition | STAT3 |
| STAT3 | localization | STAT3 |
| STAT3 | modification | STAT3 |
| STAT3 | molecular cleavage | STAT3 |
| STAT3 | phosphorylation | Akt |
| STAT3 | phosphorylation | CTNNB1 |
| STAT3 | phosphorylation | STAT3 |
| STAT3 | protein-DNA interactions | HIF1A |
| STAT3 | protein-DNA interactions | MYC |
| STAT3 | protein-DNA interactions | STAT3 |
| STAT3 | protein-DNA interactions | mir-124 |
| STAT3 | protein-protein interactions | CTNNB1 |
| STAT3 | protein-protein interactions | GSTCD |
| STAT3 | protein-protein interactions | HIF1A |
| STAT3 | protein-protein interactions | STAT3 |
| STAT3 | regulation of binding | CTNNB1 |
| STAT3 | regulation of binding | MYC |
| STAT3 | regulation of binding | STAT3 |
| STAT3 | transcription | MYC |
| STAT3 | transcription | STAT3 |
| STAT3 | translocation | STAT3 |
| STAT3 | ubiquitination | STAT3 |
| UCA1 | activation | Akt |
| UCA1 | expression | Akt |
| UCA1 | expression | GLS2 |
| UCA1 | expression | alanine |
| UCA1 | phosphorylation | Akt |
| UCA1 | protein-RNA interactions | PTBP1 |
| UCA1 | regulation of binding | PTBP1 |
| USP17L2 (includes others) | expression | MYC |
| USP17L2 (includes others) | protein-protein interactions | AKAP9 |
| USP17L2 (includes others) | protein-protein interactions | CTNNB1 |
| WNT1 | activation | Akt |
| WNT1 | activation | CTNNB1 |
| WNT1 | expression | CTNNB1 |
| WNT1 | expression | MYC |
| WNT1 | localization | CTNNB1 |
| WNT1 | localization | MYC |
| WNT1 | molecular cleavage | CTNNB1 |
| WNT1 | phosphorylation | Akt |

| | | |
|---|---|---|
| WNT1 | phosphorylation | CTNNB1 |
| WNT1 | protein-protein interactions | CTNNB1 |
| WNT1 | regulation of binding | CTNNB1 |
| WNT1 | regulation of binding | MYC |
| mir-124 | RNA-RNA interactions: microRNA targeting | STAT3 |
| mir-124 | activation | STAT3 |
| mir-124 | expression | IL2 |
| mir-124 | expression | PTBP1 |
| mir-124 | expression | STAT3 |
| mir-124 | phosphorylation | STAT3 |
| phosphatidylinositol-3,4,5-triphosphate | activation | Akt |
| phosphatidylinositol-3,4,5-triphosphate | chemical-protein interactions | Akt |
| phosphatidylinositol-3,4,5-triphosphate | localization | Akt |
| phosphatidylinositol-3,4,5-triphosphate | phosphorylation | Akt |
| phosphatidylinositol-3,4,5-triphosphate | regulation of binding | Akt |
| phosphatidylinositol-3,4,5-triphosphate | regulation of binding | phosphatidylinositol-3,4,5-triphosphate |

**Table S3: Protein – Protein interactomes of Tat in SEVs showing the proteins and their relationships.** The analysis was conducted using the STRING: functional protein association networks (https://string-db.org/). Blue text highlights key proteins present in the interactomes of Tat in SEVs.

| 7SK snRNA binding Top 5 Gene Ontology (GO) processes | | |
|---|---|---|
| **Biological Process (GO)** | **Molecular Function (GO)** | **Cellular Component (GO)** |
| Positive regulation of snRNA transcription by RNA polymerase II | 7SK snRNA binding | cyclin/CDK positive transcription elongation factor complex |
| snRNA modification | Cyclin-dependent protein serine/threonine kinase activator activity | Transcription elongation factor complex |
| Negative regulation of mRNA polyadenylation | Cyclin-dependent protein serine/threonine kinase inhibitor activity | Nucleoplasm |
| Positive regulation of protein localization to Cajal body | snRNA binding | |
| Positive regulation of DNA-templated transcription, elongation | Cyclin-dependent protein serine/threonine kinase regulator activity | |
| **mRNA Processing Top 5 Gene Ontology (GO) processes** | | |
| **Biological Process (GO)** | **Molecular Function (GO)** | **Cellular Component (GO)** |
| U2-type prespliceosome assembly | Splicing factor binding | Cleavage body |
| Positive regulation of telomerase RNA reverse transcriptase activity | BRE binding | U2 snRNP |
| Modification by virus of host mRNA processing | U5 snRNA binding | U4 snRNP |
| Targeting of mRNA for destruction involved in RNA interference | U1 snRNP binding | pICln-Sm protein complex |
| miRNA transport | snRNP binding | U5 snRNP |
| **P-TEFb complex Top 5 Gene Ontology (GO) processes** | | |
| **Biological Process (GO)** | **Molecular Function (GO)** | **Cellular Component (GO)** |
| Negative regulation of mRNA polyadenylation | 7SK snRNA binding | cyclin/CDK positive transcription elongation factor complex |
| Positive regulation of transcription elongation from RNA polymerase II promoter | RNA polymerase II CTD heptapeptide repeat kinase activity | Transcription elongation factor complex |
| Positive regulation of DNA-templated transcription, elongation | snRNA binding | |
| Transcription elongation from RNA polymerase II promoter | | |
| Regulation of mRNA processing | | |

**Table S4: Gene Ontology (GO) enrichment analysis for interactomes of Tat in SEVs.** Table shows GO terms in different networks as determined by STRING: functional protein association networks (https://string-db.org/). The table is generated with interactomes of Tat in SEVs identified from two different datasets.

| 7SK snRNA binding Top 5 Gene Ontology (GO) processes | | |
|---|---|---|
| **Reactome Pathway** | **WikiPathways** | **Subcellular localization (Compartments)** |
| Interactions of Tat with host cellular proteins | Initiation of transcription and translation elongation at the HIV-1 LTR | P-TEFb complex |
| SMAD2/SMAD3:SMAD4 heterotrimer regulates transcription | Male infertility | DSIF complex |
| Pausing and recovery of HIV elongation | | NELF complex |
| HIV elongation arrest and recovery | | |
| Formation of RNA Pol II elongation complex | | |
| Formation of HIV elongation complex in the absence of HIV tat | | |
| Tat-mediated HIV elongation arrest and recovery | | |
| RNA Polymerase II Pre-transcription Events | | |
| **mRNA Processing Top 5 Gene Ontology (GO) processes** | | |
| **Reactome Pathway** | **WikiPathways** | **Subcellular localization (Compartments)** |
| Inhibition of Host mRNA Processing and RNA Silencing | mRNA processing | U2-type prespliceosome |
| Processing of Intronless Pre-mRNAs | | Cleavage body |
| SLBP independent Processing of Histone Pre-mRNAs | | U4 snRNP |
| Processing of Capped Intronless Pre-mRNA | | U2 snRNP |
| SLBP Dependent Processing of Replication-Dependent Histone Pre-mRNAs | | pICln-Sm protein complex |
| **P-TEFb complex Top 5 Gene Ontology (GO) processes** | | |
| **Reactome Pathway** | **WikiPathways** | **Subcellular localization (Compartments)** |
| Interactions of Tat with host cellular proteins | Initiation of transcription and translation elongation at the HIV-1 LTR | P-TEFb complex |
| SMAD2/SMAD3:SMAD4 heterotrimer regulates transcription | | cyclin/CDK positive transcription elongation factor complex |
| Tat-mediated HIV elongation arrest and recovery | | DSIF complex |
| Pausing and recovery of Tat-mediated HIV elongation | | NELF complex |
| Pausing and recovery of HIV elongation | | Nucleoplasm |

**Table S5: Subcellular localization (compartments) analysis for interactomes of Tat in SEVs.** Table shows Reactome and WikiPathways Pathway identified GO processes.

| Name | Z-score | Confidence |
|---|---|---|
| *NELFE* | **6.7** | ★★★★☆ |
| *SUPT5H* | 6.6 | ★★★★☆ |
| *CDK9* | 6.6 | ★★★★☆ |
| *CCNT1* | 6.3 | ★★★★☆ |
| *NELFB* | 6.3 | ★★★★☆ |
| *SUPT4H1* | 6.1 | ★★★★☆ |
| *NELFA* | 5.8 | ★★★☆☆ |
| *HEXIM1* | 5.7 | ★★★☆☆ |
| *7SK* | 5.7 | ★★★☆☆ |
| *BRD4* | 5.6 | ★★★☆☆ |
| *RBM8A* | 5.5 | ★★★☆☆ |
| *CDK7* | 5.5 | ★★★☆☆ |
| *JAG2* | 5.5 | ★★★☆☆ |
| *PROKR2* | 5.5 | ★★★☆☆ |
| *MEPCE* | 5.4 | ★★★☆☆ |
| *AFF4* | 5.4 | ★★★☆☆ |
| *POLR2M* | 5.4 | ★★★☆☆ |
| *ELL2* | 5.1 | ★★★☆☆ |
| *TCEA2* | 5.1 | ★★★☆☆ |
| *TCEA3* | 5.1 | ★★★☆☆ |

**Table S6: Top 20 human genes for NELF complex.** The table consists of proteins in complex with NELF proteins shown in Figure 4A. [1]The subcellular localizations are derived from database annotations, automatic text mining of biomedical literature, and sequence-based predictions, and are thus not typically qualitative. The confidence of each association is signified by stars, where ★★★★★ is the highest confidence and ★☆☆☆☆ is the lowest. Developed by Janos Binder, Sune Frankild, Kalliopi Tsafou, Christian Stolte, Sean O'Donoghue, Reinhard Schneider, and Lars Juhl Jensen from the Novo Nordisk Foundation Center for Protein Research (CPR), the Luxembourg Centre for Systems Biomedicine (LCSB), and the Commonwealth Scientific and Industrial Research Organization (CSIRO).

| **NELF complex** | | |
|---|---|---|
| **Biological Process (GO)** | **Molecular Function (GO)** | **Cellular Component (GO)** |
| Negative regulation of mRNA polyadenylation | 7SK snRNA binding | NELF complex |
| Negative regulation of DNA-templated transcription elongation | Chromatin binding | cyclin/CDK positive transcription elongation factor complex |
| Negative regulation of transcription elongation from RNA polymerase II promoter | RNA binding | Transcription elongation factor complex |
| Positive regulation of viral transcription | | |
| Negative regulation of gene expression | | |

**Table S7: Gene Ontology (GO) enrichment analysis for NELF complex.** Table shows GO terms in NELF complex as determined by STRING: functional protein association networks (https://string-db.org/).

**Table S8: Interactomes of NF-κB p65 in SEVs.** The proteins were identified using mass spectrometric (MS) analysis. Data were generated from 2 biological replicates. Table is provided in .xlsx format in the online supplementary materials.

| Diseases or Functions Annotation | p-value | Associated Molecules | Regulation |
|---|---|---|---|
| Neurodevelopmental disorder with cataracts, poor growth and dysmorphic facies | 0.000168 | INTS1 | UP |
| Ciliogenesis of motile primary cilium | 0.000168 | AKAP9 | UP |
| Long QT syndrome 11 | 0.000168 | AKAP9 | UP |
| Decompaction of heterochromatin | 0.000168 | BRD2 | UP |
| Dissociation of Golgi apparatus | 0.000337 | AKAP9 | UP |
| Regulation of vascular smooth muscle cells | 0.000505 | ARHGEF28 | UP |
| Arrest in cell cycle progression of eye cell lines | 0.000505 | AKAP9 | UP |
| Lymphomagenesis of diffuse large B-cell lymphoma | 0.000505 | BRD2 | UP |
| Long QT syndrome 2 | 0.00101 | AKAP9 | UP |
| Translocation of microtubule organizing centers | 0.00118 | AKAP9 | UP |
| Autosomal dominant long-QT syndrome | 0.00168 | AKAP9 | UP |
| Inflammatory response of bone marrow-derived macrophages | 0.00168 | BRD2 | UP |
| Ploidy of cervical cancer cell lines | 0.00168 | AKAP9 | UP |
| Organization of neurofilaments | 0.00185 | ARHGEF28 | UP |
| Release of beta-estradiol | 0.00202 | BRD2 | UP |
| Long QT syndrome 1 | 0.00202 | AKAP9 | UP |
| Processing of snRNA | 0.00219 | INTS1 | UP |
| Survival of glioma cells | 0.00219 | BRD2 | UP |
| Arrest in cell cycle progression of epithelial cell lines | 0.00286 | AKAP9 | UP |
| Sudden cardiac death | 0.00303 | AKAP9 | UP |
| Localization of protein-protein complex | 0.0032 | AKAP9 | UP |
| Separation of centrosome | 0.0032 | AKAP9 | UP |
| Proliferation of embryoblast | 0.00353 | INTS1 | UP |
| Repolarization of cellular membrane | 0.00353 | AKAP9 | UP |
| Migration of eye cell lines | 0.0037 | AKAP9 | UP |
| Formation of cytoplasmic inclusions | 0.0037 | ARHGEF28 | UP |
| Duplication of centriole | 0.00421 | AKAP9 | UP |
| Proliferation of embryonic stem cell lines | 0.00471 | BRD2 | UP |
| Increased Levels of Blood Urea Nitrogen | 0.00538 | BRD2 | UP |
| Neuritogenesis of neurons | 0.00538 | ARHGEF28 | UP |
| Mitosis of eye cell lines | 0.00605 | BRD2 | UP |
| Synthesis of beta-estradiol | 0.00672 | BRD2 | UP |
| Nucleation of microtubules | 0.00739 | AKAP9 | UP |
| Endometrial cancer | 0.00784 | AKAP9, ARHGEF28, BRD2, INTS1 | UP |
| Growth of blastocyst | 0.0079 | INTS1 | UP |
| Hereditary Wolff-Parkinson-White syndrome | 0.00806 | AKAP9 | UP |
| Homologous recombination repair of bone cancer cell lines | 0.00856 | BRD2 | UP |
| Mitosis of epithelial cell lines | 0.00873 | BRD2 | UP |
| Homologous recombination repair of sarcoma cell lines | 0.00873 | BRD2 | UP |
| Fragmentation of Golgi apparatus | 0.0094 | AKAP9 | UP |
| Missegregation of chromosomes | 0.0099 | BRD2 | UP |
| Ventricular fibrillation | 0.0107 | AKAP9 | UP |
| Diffuse B-cell lymphoma | 0.0109 | ARHGEF28, BRD2 | UP |
| Cytokinesis of cervical cancer cell lines | 0.0112 | AKAP9 | UP |
| Hepatobiliary carcinoma | 0.0116 | AKAP9, ARHGEF28, BRD2, INTS1 | UP |
| Binding of GTP | 0.0116 | ARHGEF28 | UP |
| Formation of nucleosomes | 0.0116 | BRD2 | UP |
| Autosomal recessive cataract disease | 0.0119 | INTS1 | UP |
| Concentration of beta-estradiol | 0.0127 | BRD2 | UP |
| Hepatobiliary system cancer | 0.0132 | AKAP9, ARHGEF28, BRD2, INTS1 | UP |
| Polymerization of microtubules | 0.0141 | AKAP9 | UP |
| Sensitivity of breast cancer cell lines | 0.0151 | BRD2 | UP |
| Ventricular tachycardia | 0.0152 | AKAP9 | UP |

| | | | |
|---|---|---|---|
| Generation of embryonic cell lines | 0.0161 | AKAP9 | UP |
| Differentiation of embryonic stem cell lines | 0.0162 | BRD2 | UP |
| Growth of organism | 0.0177 | ARHGEF28, INTS1 | UP |
| Movement of adenocarcinoma cell lines | 0.0181 | ARHGEF28 | UP |
| Colorectal cancer | 0.0193 | AKAP9, ARHGEF28, BRD2, INTS1 | UP |
| Primary dilated cardiomyopathy | 0.0214 | AKAP9 | UP |
| Basal cell carcinoma | 0.0215 | AKAP9, ARHGEF28, INTS1 | UP |
| Quantity of ovarian follicle | 0.0222 | BRD2 | UP |
| Organismal death | 0.0236 | AKAP9, BRD2, INTS1 | UP |
| Clear-cell ovarian carcinoma | 0.0249 | AKAP9 | UP |
| Migration of epithelial cell lines | 0.0252 | AKAP9 | UP |
| Closure of neural tube | 0.0255 | BRD2 | UP |
| Development of fibroblast cell lines | 0.026 | AKAP9 | UP |
| Lymphoreticular neoplasm | 0.0264 | AKAP9, ARHGEF28, BRD2 | UP |
| Segregation of chromosomes | 0.0267 | BRD2 | UP |
| Development of genitourinary system | 0.0282 | AKAP9, BRD2 | UP |
| Cell cycle progression | 0.0299 | AKAP9, BRD2 | UP |
| Arrest in growth of embryo | 0.0328 | INTS1 | UP |
| Mature B cell malignant tumor | 0.033 | ARHGEF28, BRD2 | UP |
| Formation of cellular protrusions | 0.0342 | AKAP9, ARHGEF28 | UP |
| Hypertrophic cardiomyopathy | 0.0351 | AKAP9 | UP |
| Lifespan of organism | 0.0356 | BRD2 | UP |
| Renal clear cell adenocarcinoma | 0.0374 | AKAP9, INTS1 | UP |
| Serine phosphorylation of peptide | 0.0388 | AKAP9 | UP |
| Migration of fibroblasts | 0.039 | ARHGEF28 | UP |
| Autosomal recessive mental retardation | 0.0416 | INTS1 | UP |
| Pharyngeal squamous cell carcinoma | 0.0437 | AKAP9 | UP |
| Mixed neoplasia | 0.0492 | ARHGEF28, BRD2 | UP |

**Table S9: Diseases or Functions Annotation of interactomes of Tat and NF-κB p65 in SEVs.** The analysis was conducted using Ingenuity Pathway Analysis (IPA). Blue text highlights functional annotations related to decompaction of heterochromatin and inflammatory response of bone marrow-derived macrophages mediated by BRD2 that are of interest.

| GO Name | Raw P-value | Adjusted P-value |
|---|---|---|
| Regulation of metabolic process | 0.0001 | 0.7766 |
| Epithelial cell morphogenesis | 0.0002 | 0.7766 |
| Regulation of gene silencing by miRNA | 0.0006 | 0.7766 |
| Regulation of post-transcriptional gene silencing | 0.0007 | 0.7766 |
| Regulation of gene silencing by RNA | 0.0007 | 0.7766 |
| Mesoderm migration involved in gastrulation | 0.0007 | 0.7766 |
| Male somatic sex determination | 0.0007 | 0.7766 |
| Prostate induction | 0.0007 | 0.7766 |
| Prostate field specification | 0.0007 | 0.7766 |
| Activation of prostate induction by androgen receptor signaling pathway | 0.0007 | 0.7766 |

**Table S10: Top 10 enriched Gene Ontology (GO) categories for BRD2 interactors.** Analysis was conducted by querying functional database (PPI BIOGRID) for Network Topology-based Analysis (NTA) using WebGestalt (WEB-based GEne SeT AnaLysis Toolkit, https://www.webgestalt.org/). Both Raw and Adjusted P-values were determined by the software using inbuilt algorithm.

**Data file S1: All proteins from SEVs identified in complex with NF-κB.** Pooled uninfected subjects (HIV-ART-) (n=15, split into 2 groups of 7 for group 1 and 8 for group 2) were used for the study. The proteins were identified using mass spectrometric (MS) analysis. Data were generated from 2 biological replicates. Data are provided in an Excel sheet in the online supplementary materials.

**Data file S2: All proteins from SEVs identified in complex with Tat**. Pooled uninfected subjects (HIV-ART-) (n=15, split into 2 groups of 7 for group 1 and 8 for group 2) were used for the study. The proteins were identified using mass spectrometric (MS) analysis. Data were generated from 2 independent experiments. Data are provided in an Excel sheet in the online supplementary materials.
